# An alternatively spliced zebrafish *jnk1a* transcript has an essential and non-redundant role in development of the first heart field derived proximal ventricular chamber

**DOI:** 10.1101/546184

**Authors:** A Santos-Ledo, S Washer, T Dhanaseelan, P Chrystal, T Papoutsi, DJ Henderson, B Chaudhry

## Abstract

Alternative splicing is a ubiquitous mechanism for producing different mRNA species from a single gene, resulting in proteomic diversity. Despite potential for regulating embryogenesis, its developmental role remains under-investigated. The *Jun kinase* (*Jnk*) genes, considered downstream effectors of the non-canonical Wnt planar cell polarity pathway, utilise extensive and evolutionarily-conserved alternative splicing. Although many PCP members are associated with heart malformation, the role of *Jnk* genes in cardiac development, and specifically which alternatively spliced transcripts orchestrate these processes, remain unknown. In this study we exploit the *jnk1* duplication and subspecialisation found in zebrafish to reveal an essential and non-redundant requirement for *jnk1a* in cardiac development. We characterise alternatively spliced *jnk1a/jnk1b* transcripts and demonstrate that hypoplasia of the proximal ventricular component, which corresponds to human hypoplastic left ventricle, can only be rescued by the *jnk1a Ex7 Lg* transcript. These studies highlight the importance of Jnk signalling and alternative splicing in heart development

## Introduction

Over 1% of the population have structural congenital heart disease (CHD) [1]. Despite several studies finding genes associated with syndromic heart disease [2-5], genes contributing to CHD in the vast majority of patients remain elusive. An alternative to population based genomic studies is the use of developmental biology approaches to understand how the heart forms and identify pathways and genes that are relevant to CHD. One genetic pathway that has been shown to be important in pre-clinical models is the non-canonical Wnt, planar-cell polarity (PCP) pathway. Disruption of key genes such as *Vangl2, Dvl2* and *Celsr1*, reviewed in [6], have been shown to cause cardiac malformations, specifically involving addition of second heart field precursors [7]. Inactivation of downstream pathway effectors, such as Rac1 [8,9] and Rho Kinase (ROCK) 1 and 2 [10] also produce cardiac defects.

The c-Jun N-terminal kinases (JNK1, 2 and 3) are members of the mitogen activated protein kinase (MAPK) family, but are also recognised as mediators of PCP signalling, acting downstream of RhoA. These stress-responsive serine/threonine kinases are implicated in a diverse range of biological processes including embryonic development, brain signalling, tumour survival and metastasis, through control of cell proliferation, cell death and direct regulation of gene expression (reviewed in [11]). Jnk is phosphorylated by MAP kinase kinases (MAP2K), which in turn are activated by MAP kinase kinase kinases (MAP3K) [12], but also integrates signals from the RAS and AKT pathways, for example, the R497Q mutation in *SOS1*, found in patients with Noonan syndrome, has been shown to directly activate Jnk [13]. Alternative splicing is a universally applied gene regulatory mechanism that increases the diversity of gene products; up to 95% of all human genes undergo alternative splicing (14). The *Jnk* genes have strong evolutionary conservation and undergo alternative splicing to provide a range of transcripts. For each gene, alternative usage of two neighbouring exons creates alternative peptide structures. In addition, *Jnk1* and *Jnk2,* but not *Jnk3*, have truncated or extended C termini, which are manifest as either 46 or 54 kDa-sized peptides. An alternative translation initiation site for *Jnk3* increases the range of transcripts to 10 for that gene [15, 16]. The functional significance of these different transcripts is unknown. Differential expression of different splice isoforms of *Jnk1* has been shown within NIH3T3 fibroblasts [17], but expression of the alternatively spliced transcripts has not been examined in animal studies. Little is known of their roles in heart development [11]. Null mutant mice are available and whilst both *Jnk1* and *Jnk2* are co-expressed in almost all cells and tissues, *Jnk3* has been said to be expressed predominantly in the brain, testes and heart [18]. This redundancy of expression and likely function is one reason why identifying the roles of *JNK* has been difficult, particularly those in cardiovascular development. The *Jnk1* null mouse is viable and fertile, with no obvious malformations, but has abnormal differentiation of T-helper cells and reduced adipose tissue [18]. Similarly, the *Jnk2* null mouse also has abnormalities of T-cell function and abnormal brain development [18, 19, 20]. In contrast the *Jnk3* null mouse is essentially normal and despite the specific expression of *Jnk3* within the central nervous system, has normal brain histology [18, 19]. Attempts to unravel these roles in compound null mutants have met limited success. Mice null for both *Jnk1* and *Jnk2* die by embryonic day E11, with evidence of extensive apoptosis in the brain. No specific cardiac abnormalities were noted although some embryos had cardiac dilatation, interpreted to be a non-specific finding [19] although the reason for this interpretation is unclear.

The zebrafish has become established as an important pre-clinical model, especially in developmental studies, because of the high degree of conservation of vertebrate genes and developmental processes. The unique combination of transgenic reporter lines, transparent embryos, and ease of genetic manipulation through morpholino and CRISPR-Cas9 genome editing are practical advantages over other laboratory animals [21]. Zebrafish are particularly useful for cardiovascular developmental studies as they can survive without a functional circulation for up to one week [22]. Although the zebrafish has only a single atrium and ventricle, the initial fundamental processes of vertebrate heart formation are conserved. An initial heart tube forms from first heart field (FHF) progenitors and further addition of second heart field (SHF) progenitors augment the ventricular mass and atrium [21]. Thus, although not septated, the zebrafish ventricle contains a proximal FHF derived portion that is equivalent to the FHF derived left ventricle and a distal SHF derived component which is the equivalent to the right ventricle [23, 24]. A complicating factor in zebrafish genetics is a genome duplication event [25], such that some genes persist as duplicated paralogs, often with sub-functionalisation and different expression patterns. For example, there is only one persisting functional gene for *jnk2* and *jnk3*, but *jnk1* remains duplicated as *jnk1a* and *jnk1b*.

In this study we have uncovered the role of *jnk1* in heart development using zebrafish to overcome embryonic death which has limited mouse studies. These studies show developmental stage and tissue level changes in *jnk1* transcripts brought about by differential gene expression and alternative splicing. Using both CRISPR null mutants and morpholinos we show a specific requirement for *jnk1a* in development of the FHF-derived ventricle; the evolutionary equivalent of the left ventricle in man. Moreover, we show that only one specific evolutionarily conserved alternative splice transcript is responsible for this role and that its production is associated only with *jnk1a*, as a result of sub-specialisation.

## Results

### Bioinformatic analysis of the duplicated zebrafish genome suggests sub-functionalisation of jnk1

We used the zebrafish to understand the developmental roles of *jnk* genes in vertebrate cardiac morphogenesis. Initial bioinformatic analysis of the zebrafish genome using Ensembl (assembly GRCz11) [26] indicated that the zebrafish *jnk1a* and *jnk1b* genes are paralogs and orthologs of human *JNK1* (Figure 1A, B). Expression studies in mice [19,20] show that *Jnk2* is widely and strongly expressed throughout the developing embryo and *Jnk3* is reported to be expressed in the heart. Before concentrating on jnk1 function we examined the expression of *jnk2* and *jnk3* in embryonic development. We noted a similar broad expression pattern of *jnk2* in developing zebrafish as described in mouse (Figure 1C). Expression of *jnk3* was limited to the developing central nervous system at 24 hours post fertilisation (hpf) and we were not able to show *jnk3* expression in the developing zebrafish heart by wholemount *in-situ* hybridisation (WISH) (Figure 1C). These findings were confirmed by RT-PCR of extracted hearts, with *jnk3* transcripts clearly detected at 48 hpf but only weak expression of *jnk3* detected at 24 hpf. *jnk2* was detected at both time points (Figure 1D). We therefore focussed our attention on *jnk1a* and *jnk1b* expression and in particular wanted to look at expression of alternatively spliced transcript variants Expression of *jnk1* has been reported in zebrafish [27] but at that time the gene duplication was not recognised, and the presence of alternative spliced transcripts not attempted. In order to examine the spatial-temporal expression of these *jnk1* paralogues, we firstly sub-cloned *jnk1a* and *jnk1b* transcripts from pooled whole embryos at 24, 48 and 72 hpf and analysed their sequence, revealing conservation of the alternative splicing patterns seen in human *JNK1*. Each gene revealed four transcript variants indicating alternative exon 7/8 usage and an alternative splice re-entry site encoding short (Sh) and long (Lg) C-terminal peptides (Figure 1E and sequences in Supplementary Table 2). The splicing events required to produce these transcripts correspond to exon 6a/6b alternative splicing and the p46 or p54 C-terminal extension variants of human *JNK1* [11]. Comparison of the predicted zebrafish jnk1a/jnk1b peptides with human JNK1 peptides [28] suggested the potential for sub-functionalisation. The zebrafish *jnk1a* gene, but not the *jnk1b*, was capable of producing Ex7 Lg and Ex7 Sh products, equivalent to human *JNK1* Ex6a p46 and p54 C-terminal extension variants. In complementary fashion, *jnk1b* appeared capable of producing an Ex8 Sh peptide comparable with the human *JNK1* Ex6b p46 variant. However, neither *jnk1* zebrafish gene produced a product typical of the human *JNK1* Ex6b p54 splice variant. Specifically, the exon 8 peptide of the zebrafish *jnk1a* was not well conserved in comparison with human *JNK1* exon 6b peptide and, although the *jnk1b* Ex8 peptide was conserved, the C-terminal peptide extension provided by *jnk1b* was extremely divergent from the human C-terminal extension (Figure 1F).

**Figure 1.**
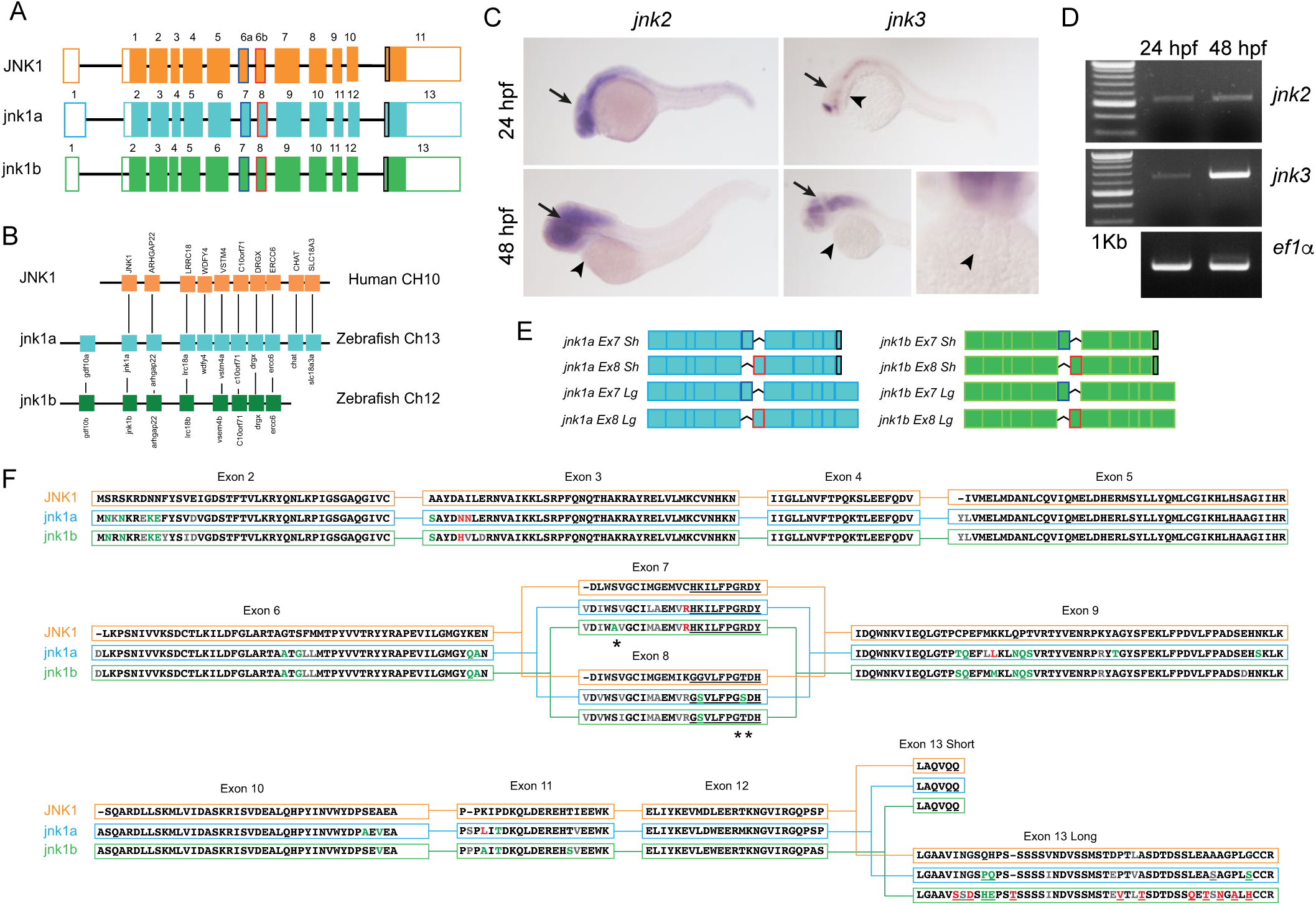
Human *JNK1* is conserved in zebrafish, although duplicated as *jnk1a* and *jnk1b*. (**A**) Despite different exon numbering i.e. 6a/6b in human and 7/8 in zebrafish orthologues, exon structure is entirely conserved. (**B**) *jnk1a, jnk1b* and *JNK1* lie in syntenic chromosomal regions. (**C**) WISH demonstrated expression of *jnk2* and *jnk3* in the developing brain (arrows) but not heart (arrowheads). (**D**) RT-PCR of extracted hearts indicates late onset of *jnk3* gene expression. (**E**) Cartoon showing all zebrafish *jnk1a/b* transcripts resulting from alternate exon 7/8 usage and differing C-terminal extension. (**F**) Translation of zebrafish *jnk1a* and *jnk1b* transcripts indicates conservation and subspecialisation of zebrafish jnk1a and jnk1b with human JNK1 peptides when allowance for favourable (green text) or neutral (grey text) amino acid substitutions [38]. Amino acids coded by exon 7 within *jnk1a* match those in human exon 6a, but *jnk1b* lacks a conserved serine residue (*). In reciprocal fashion, exon 8 in *jnk1b* encodes a conserved threonine residue seen in *JNK1* exon 6b, whilst *jnk1a* codes for serine in exon 8 (**). The short C-terminal peptide sequence was completely conserved between the species. The C-terminal extension seen in jnk1a was almost identical to the human, but the jnk1b extension differed by 9/39 amino acids including insertion of an additional threonine residue.

### Specific jnk1a splice variants may play a role in early ventricular morphogenesis

To understand the implications of the *jnk1a-jnk1b* gene duplication event we attempted a spatiotemporal analysis of all eight transcripts by WISH. Riboprobes based on the full-length transcripts were used in high-stringency conditions. As expected, high levels of *jnk1a* and *jnk1b* transcripts were expressed throughout the developing brain. Despite the degree of similarity between each of the gene products, it was possible to identify different expression patterns corresponding to the different transcripts, particularly in the developing heart (Figure 2A and Supplementary Figure 1). At 24 hpf: *jnk1a Ex8 Lg*; *jnk1b Ex8 Sh;* and *jnk1b Ex8 Lg* were expressed in the heart as well as throughout the developing head structures (Figure 2 B-D). One specific *jnk1a* splice variant, *jnk1a Ex7 Lg*, was strongly and specifically expressed solely in the developing heart at 24 hpf (Figure 2A). By 48 hpf this transcript was clearly localised to the proximal, rather than distal, part of the developing ventricular chamber, with some localised expression in the adjacent part of the developing atrium.

**Figure 2.**
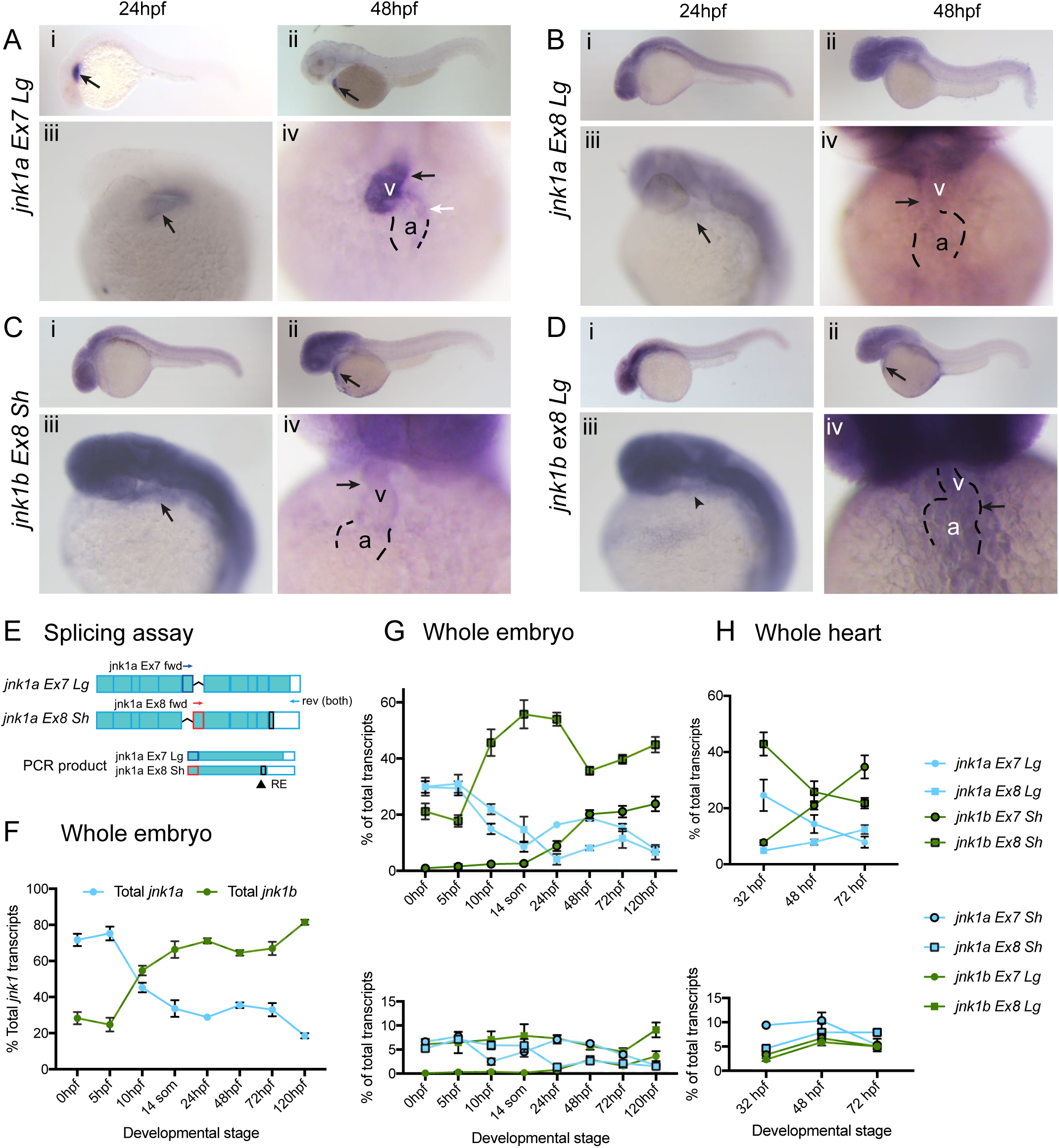
Expression of *jnk1a/b* transcripts in development. (**A-D**) Expression patterns of the four *jnk1* transcripts most highly expressed in the heart (arrows) at 24hfp (i, iii), 48hpf (ii, iv) from lateral (i, ii) and left oblique views (iii, iv). (**A**) *jnk1a Ex7 Lg* is expressed in the cardiac cone (arrow) at 24hpf. At 48hpf strong expression is seen in the proximal part of the cardiac ventricle (v; black arrow) and weak expression in some cells in the outer curvature of the atrium (a; white arrow). (**B**) *jnk1a Ex8 Lg* is expressed at low level in in both the atrium and ventricle. (**C**) At 24hpf *jnk1b ex8 Sh* expression in the cardiac cone is difficult to determine due to expression in overlapping head structures (**C i, III**), but is clearly seen in the outflow tract (arrow), and lower level in the ventricle, by 48hpf (**C ii-iv**). (**D**) *jnk1b Ex8 Lg* is expressed in the region of the heart but is also expressed in surrounding tissues. (**E-H**) Semi-quantitative *jnk1a/b* transcript assay. (**E**) Schematic of assay for *jnk1a* (see methods for full details). RNA is extracted and a cDNA pool produced. PCR within linear phase performed using forward (fwd) primers specific for *jnk1a*/*1b* exons 7/8 and common reverse (rev) primers. The fraction with C-terminal extension is determined by restriction enzyme (RE) digestion. (**F**) Percentage of total *jnk1a* and *jnk1b* transcripts during embryonic development. 14 ss = 16 hpf. (**G**) Proportion of each individual alternatively spliced transcript during development in whole embryo. (**H**) Proportion of each individual transcript within isolated whole heart during development. *jnk1a* = blue, j*nk1b* = green, Ex7 = circle, Ex8 = square, black outline = Sh and no outline = Lg.

### Expression profile of jnk transcripts different between heart and whole embryo

To confirm this expression and provide quantitative data, we designed a semi-quantitative RT-PCR/restriction enzyme assay to measure the relative expression of the individual *jnk1a* and *jnk1b* alternatively spliced transcripts (Figure 2E). This was based on RT-PCR to identify exon 7/8 usage and then a restriction digest to determine the fraction of each PCR product relating to short or long C-terminal extension (Figure 2B and Supplementary Figure 2 for validation of assay) [29]. Between fertilisation and the onset of gastrulation, 80% of *jnk1* transcripts originated from the *jnk1a* gene. However, after this *jnk1b* transcripts became increasingly abundant and by 120 hpf, 80% of *jnk1* transcript originated from *jnk1b* (Figure 2F). Throughout this phase of development, the majority of *jnk1a* transcripts encoded for the long C-terminal form with equal representation of exons 7 and 8 containing transcripts (Figure 2G). In contrast, the majority of *jnk1b* transcripts contained exon 8 and were of the short C-terminal form. These data are dominated by the forming brain and somites as major sites of *jnk1* expression in the early embryo. To understand which transcripts are specifically important in heart development we isolated whole hearts [31] at key embryonic stages. The transcript assay was performed on hearts extracted at 32 hpf, when the linear heart tube has formed from first heart field (FHF) precursors, but before significant addition of the second heart field (SHF) cells [23,32]. This was compared with 48 hpf and 72 hpf, timepoints when addition of the second heart field is complete [23,32]. The proportion of transcripts represented by the *jnk1a* gene within the heart paralleled the changes observed in the whole embryo, with falling levels of *jnk1a ex7 Lg*, and lower, but maintained, levels of *jnk1a ex8 Lg* (Figure 2H). In contrast, whilst *jnk1b Ex8 Sh* had become the most abundant transcript in the entire embryo, its levels gradually fell within the heart and by 72 hpf, *jnk1b Ex7 Sh* was the most abundant transcript.

### jnk1a and jnk1b mutants appear overtly normal

To understand the functions of *jnk1* in heart development we designed translation-blocking morpholino oligonucleotides to knock down production of jnk1a and jnk1b protein and created *jnk1a* and *jnk1b* null mutants using CRISPR-Cas9 (Figure 3 A, B). Although *jnk1a* morphants appeared normal, approximately 50% of *jnk1b* morphants exhibited curling of the tail and a similar percentage of *jnk1a/jnk1b* morphants exhibited minor pericardial oedema. Experiments were carried out using 1ng *jnk1b* MO and 2ng *jnk1a* MO as these were the minimal dosages that produced the abnormalities later identified in the mutant lines (see below), without any evidence of off target effects. For the mutants, after initial outcrossing, heterozygotes were in-crossed and resulted in zygotic (Z) null mutants that were fertile and of normal appearance. These could then be in-crossed and maintained as homozygous maternal zygotic (MZ) null lines, where eggs contained no maternally deposited *jnk* mRNA. Null mutations were confirmed both by sequencing genomic DNA (Figure 3A, B) and demonstration of nonsense mediated decay by RT-PCR (Figure 3C) in MZ null mutants. *MZjnk1a, MZjnk1b* and *MZjnk1a*/*MZjnk1b* zebrafish appeared grossly normal with normal behaviour and fertility (Figure 3D).

**Figure 3.**
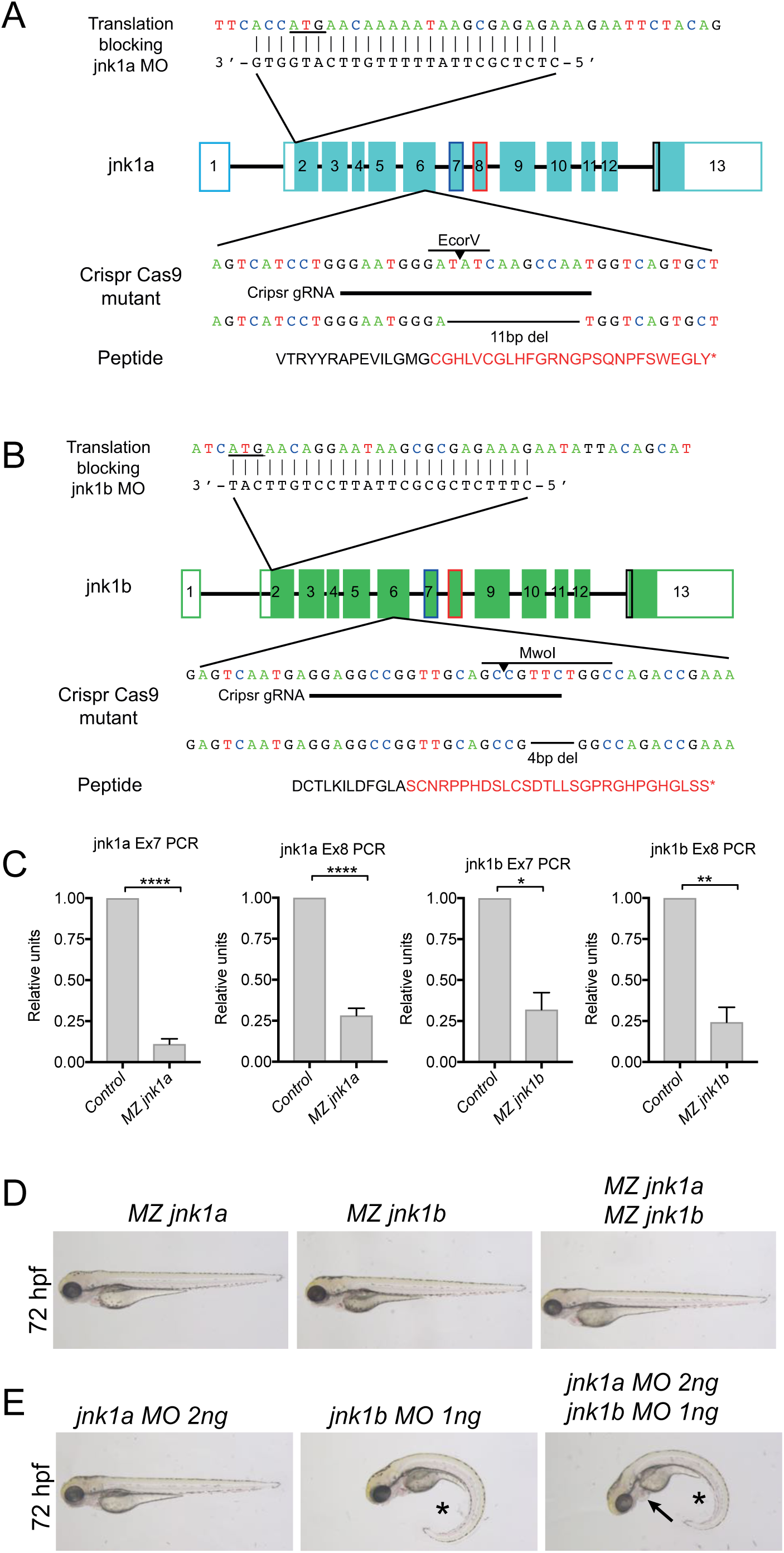
Rationale for *jnk1* mutants and morphants. Sequences of genomic DNA for *jnk1a* **(A)** and *jnk1b* **(B)** showing: relative position of translational blocking morpholino oligonucleotides (ATG underlined); targeting sequence for CRISPR-Cas9 guide RNA; deletions produced; translated peptide sequence (* = stop); and restriction enzymes that identify mutants. Both *jnk1a* and *jnk1b* mutations disrupt sequence within exon 6 and lead to loss of all four alternatively spliced transcripts. (**C**) RT-PCR using primers specific for *jnk1a* and *jnk1b* exon 7 and 8 demonstrate nonsense-mediated decay. * p<0.05 ** p<0.02 ****p<0.0001. **(D**) M*aternal zygotic (MZ) jnk1a, MZjnk1b* and *MZjnk1a/ MZjnk1b* mutants all appear grossly normal and are fertile. (**E**) *jnk1a* 2ng morphant appears normal; approximately 50% of *jnk1b* 1ng morphant displays tail curling (*) and some *jnk1a* 2ng*/jnk1b* 1ng morphants additionally display mild pericardial oedema (arrow).

### Hypoplastic first heart field ventricular segment

The highly specific expression pattern of *jnk1a Ex7 Lg* (Figure 2A) led us to examine heart development in more detail. The early stages of vertebrate heart development are conserved, including: formation of the initial heart tube from FHF cardiomyocytes; asymmetrical remodelling; and addition of cardiomyocytes derived from the SHF. Formation of the initial heart tube by migrating FHF cardiomyocytes is complete by 28-30 hpf. Immunolabeling of the heart chambers at that time indicated that the ventricular chamber was reduced in size in *MZjnk1a* mutants and 2ng *jnk1a* morphants at 28 hpf. However, there was no apparent change in *MZjnk1b* mutants and *jnk1b* morphants and furthermore no additional size reduction in *MZjnk1a*/*MZjnk1b* double mutants (Figure 4A).

**Figure 4.**
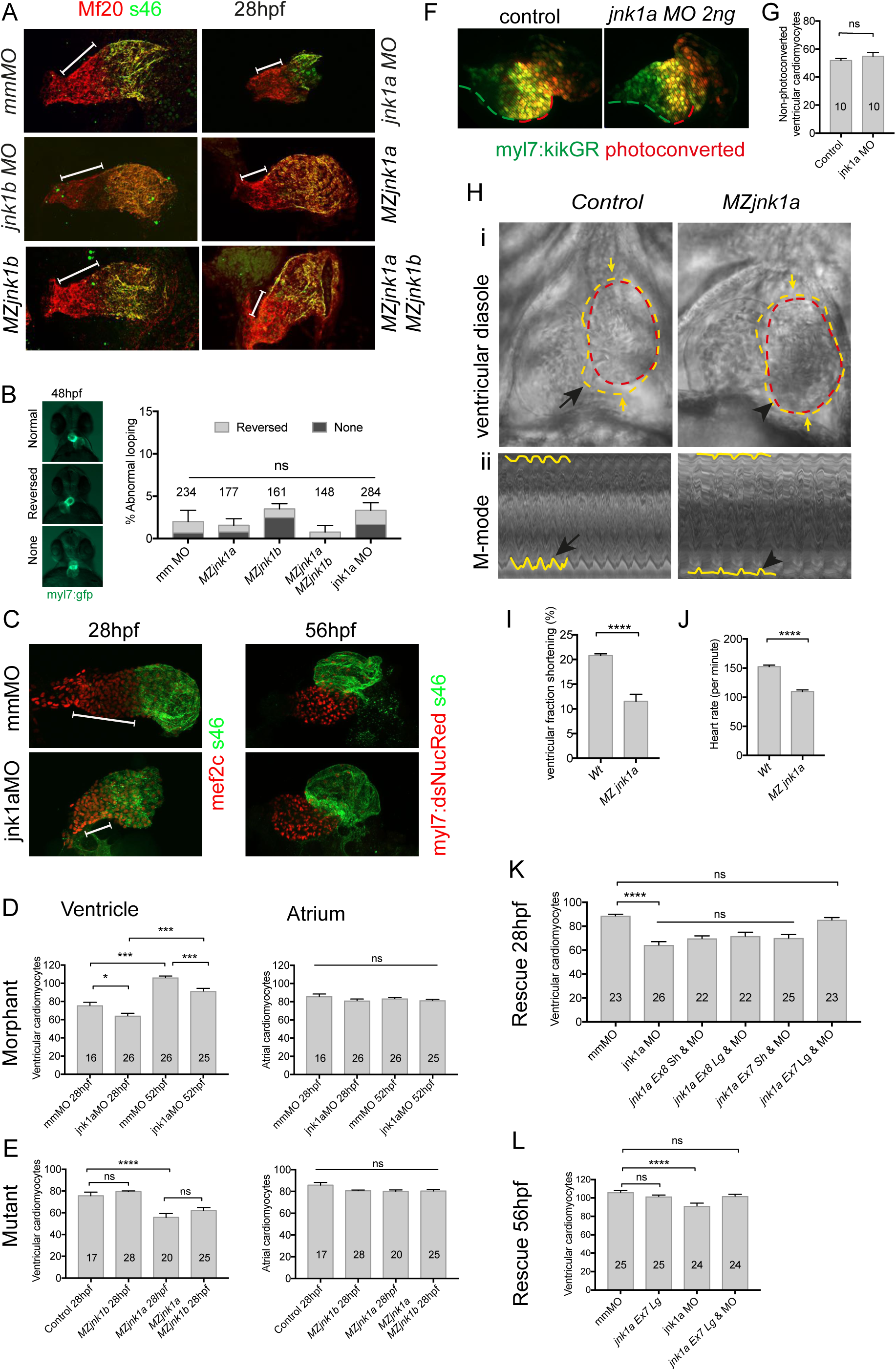
Ventricular hypoplasia in *jnk1a* knockdown. (**A**) Cardiac morphology at 28 hpf. All ventricular cardiomyocytes are labelled by MF20 antibody (red) and atrial cells by both MF20 and S46 antibody (appear yellow). Normal appearances seen in embryos exposed to 2ng mismatch morpholino (mmMO). Ventricular hypoplasia (shown by size of white bar) is seen in embryos injected with 2ng *jnk1a* morpholino, but not in those treated with *1*ng *jnk1b* morpholino. Maternal zygotic (MZ) *jnk1a* null mutants have a comparable degree of ventricular hypoplasia, which is not seen in *MZjnk1b.* There is no additional ventricular hypoplasia in *MZjnk1a/MZjnk1b* double mutants. **(B)** Minimal levels of disordered cardiac looping in *MZjnk1a* and *MZjnk1b* mutants, alone and in combination. There is no increase in cardiac looping abnormalities in *jnk1a* 2ng MO treated embryos. **(C)** Quantification of cardiomyocytes within the developing heart. Embryos at 28 hpf treated with 2ng mmMO or 2ng *jnk1a* MO immunolabelled with mef2 antibody which identifies all cardiomyocyte nuclei at this stage, and S46 antibody which identifies atrial cells. Hypoplasia of the FHF-derived ventricular segment is obvious at 28 hpf (line). (**D,E**) In embryos treated with 2ng *jnk1a* MO there is a reduction in ventricular cardiomyocytes at 28 hpf before, and at 52 hpf after, completion of the SHF cardiomyocyte accretion. Although there are increased numbers of cardiomyocytes at 56 hpf, this difference remains. There is no change in the numbers of atrial cardiomyocytes. The same reduction in FHF-derived ventricular cardiomyocytes is seen in *MZjnk1a* mutants at 28 hpf but not in *MZjnk1b* mutants and there is no change in severity of this phenotype when *jnk1a* and *jnk1b* are both absent. (**F**) Confirmation of normal SHF ventricular cardiomyocyte addition at 56hpf. The green dashed line indicates the extent of the SHF-derived ventricular component whereas the red dashed line indicates the extent of the FHF-derived ventricular component. (**G**) numbers of SHF ventricular cardiomyocytes (**H**) i) Cardiac contractile function in *MZjnk1a* mutants. Images from movies (see supplementary data) with the external margin of the ventricle identified in diastole (yellow dashed line) and systole (red dashed line). Ventricular contraction is impaired in the FHF-derived ventricular segment (black arrowhead). Ii) “M-mode” representations of the movies are obtained by resampling between the yellow arrows. Hypokinesis in the *MZjnk1a* heart (black arrowhead) can be seen, in comparison to the normal waveform of contraction in the control heart (black arrow). (**I**) Reduced fractional shortening and (**J**) heart rate in *MZjnk1a* mutants. (**K**) Only *jnk1a Ex7 Lg* mRNA was capable of rescuing the effects of *jnk1a* morpholino in reducing the FHF ventricular cardiomyocyte numbers. (**L**) *jnk Ex7 Lg mRNA* does not increase the complement of total FHF and SHF ventricular cardiomyocytes but is able to reverse the effects of *jnk1a MO* at 56hpf. * p<0.05, ** p<0.02, ***p<0.001, ****p<0.0001, ns = not significant.

The first morphological evidence of left-right asymmetrical remodelling in vertebrates is manifest as jogging of the heart tube. In most normal zebrafish embryos, the heart tube extends from the midline towards the left side of the body, whereas in 1-2% of wildtype embryos this jogging is reversed or hearts lie in the midline. Then, looping of the heart tube places the atrium to the left side of the ventricle but again, in 1-2% of normal embryos this is reversed, or the chambers remain orientated in the midline. In the *MZjnk1a, MZjnk1b, MZjnk1a*/*MZjnk1b* mutants and *jnk1a* morphants, there was no significant difference in looping disturbance at 48 hpf (Figure 4B). However, a small degree of jogging and looping disturbance was noted in *jnk1b* morphants at the same stage; further analyses regarding the role of *jnk* in left-right patterning is described elsewhere (Santos-Ledo et al; in preparation). Following formation of the initial heart tube and concomitant with asymmetrical remodelling is the continued addition of SHF cells to the heart tube. In birds and mammals with septated hearts SHF cells form the right ventricle, but in the zebrafish, which maintains a single unseptated ventricle, the SHF cells add to the distal part of the common ventricle [23,24,32]. To fully assess the size of the FHF and SHF atrial ventricular components we counted the number of atrial and ventricular cardiomyocytes within the heart at 28 hpf, when the FHF cells have added but the SHF cells have not, and at 56 hpf when the SHF accretion is complete but growth by replication of CMCs has not yet begun. In 2ng *jnk1a* morphants there was a 15% reduction in the number of ventricular cardiomyocytes within the heart at 28 hpf (Figure 4 C, D). At 56 hpf, the number of CMCs was increased equally in both control and *jnk1a* morphant ventricles indicating that the SHF accretion was unaffected by *jnk1a* knockdown, and the reduction in ventricular size was less apparent (Figure 4C, D). Analysis of *MZjnk1a* mutants at the same stages indicated a 26% reduction in ventricular cardiomyocytes at 28 hpf. There was no significant reduction in *MZjnk1b* mutants and no additional reduction in *MZjnk1a*/*MZjnk1b* compared to *MZjnk1a* mutants (Figure 4E). To confirm that there was normal addition of SHF progenitors to the heart tube we used a zebrafish line expressing the photoconvertible kikGR reporter under the control of the *myh7* promoter [23]. Photo-conversion of kikGR at 20 hpf identifies FHF cardiomyocytes, in which the *myh7* promoter is already active, and allows them to be distinguished from SHF CMCs that only later activate the *myh7* promoter and thus only exhibit non-photoconverted protein [23]. Analysis at 50 hpf showed that here was no difference between *jnk1a* morpholino treated embryos and controls (Figure 4 F,G).

We asked if there might be functional consequences from this significant and localised deficiency of FHF ventricular cardiomyocytes. We therefore examined heart function at 72 hpf in control and *MZjnk1a* mutant embryos. Using video microscopy it was evident that the FHF ventricular component, adjacent to the atrioventricular valve, was of reduced size and hypokinetic in the *MZjnk1a* mutant compared with their control counterparts (Figure 4H) leading to a reduction in ventricular fractional shortening (Figure 4I). There was also a reduction in heart rate in the mutant embryos (Figure 4J).

In order to confirm that the defects in formation of the FHF ventricular component were specific to *jnk1a* knockdown, and to establish whether they related specifically to any of the isoforms, the four *jnk1a* transcripts: *Ex7 Sh*; *Ex7 Lg*; *Ex8 Sh;* and *Ex8 Lg* were evaluated with regard to their ability to rescue the ventricular cardiomyocyte deficit at 28 hpf (Figure 4H). One cell-stage embryos were injected simultaneously with the *jnk1a* morpholino and 100pg capped mRNA corresponding to the individual transcripts. Only the *jnk1a Ex7 Lg* transcript was able to fully rescue the FHF cardiomyocyte deficit and the effect was still evident at 56 hpf (Figure 4I). None of the other three *jnk1a* transcripts, separately or in combination, were able to do this. These data suggest that *jnk1a Ex7 Lg* is specifically required to produce the normal FHF derived ventricular component, the equivalent of the left ventricle in septated hearts.

### Mechanism of FHF ventricular cardiomyocyte deficiency

To understand how disruption of *jnk1a* function might lead to this specific deficiency of FHF ventricular cardiomyocytes, we followed the development of cardiomyocytes from their appearance in the anterior lateral plate mesoderm (ALPM) to their coalescence to the primary heart tube. At the 8-somite stage (ss) cells in the parasagittal ALPM express the key transcriptional regulators *nkx2.5, hand2* and *gata5*. In *jnk1a* morphants the expression patterns of these genes all appeared normal (Figure 5A). Whilst the expression pattern of these genes includes other non-cardiomyocyte cells, ventricular cardiomyocytes could be specifically detected by ventricular specific myosin (*myh7*) expression at 15ss (Figure 5B). At this point there was no difference between the expression patterns of *jnk1a* morphants and controls. However, at 18ss the expression pattern of *myh7* revealed midline convergence of ventricular cardiomyocytes occurring in control embryos, but not *jnk1a* morphants.

**Figure 5.**
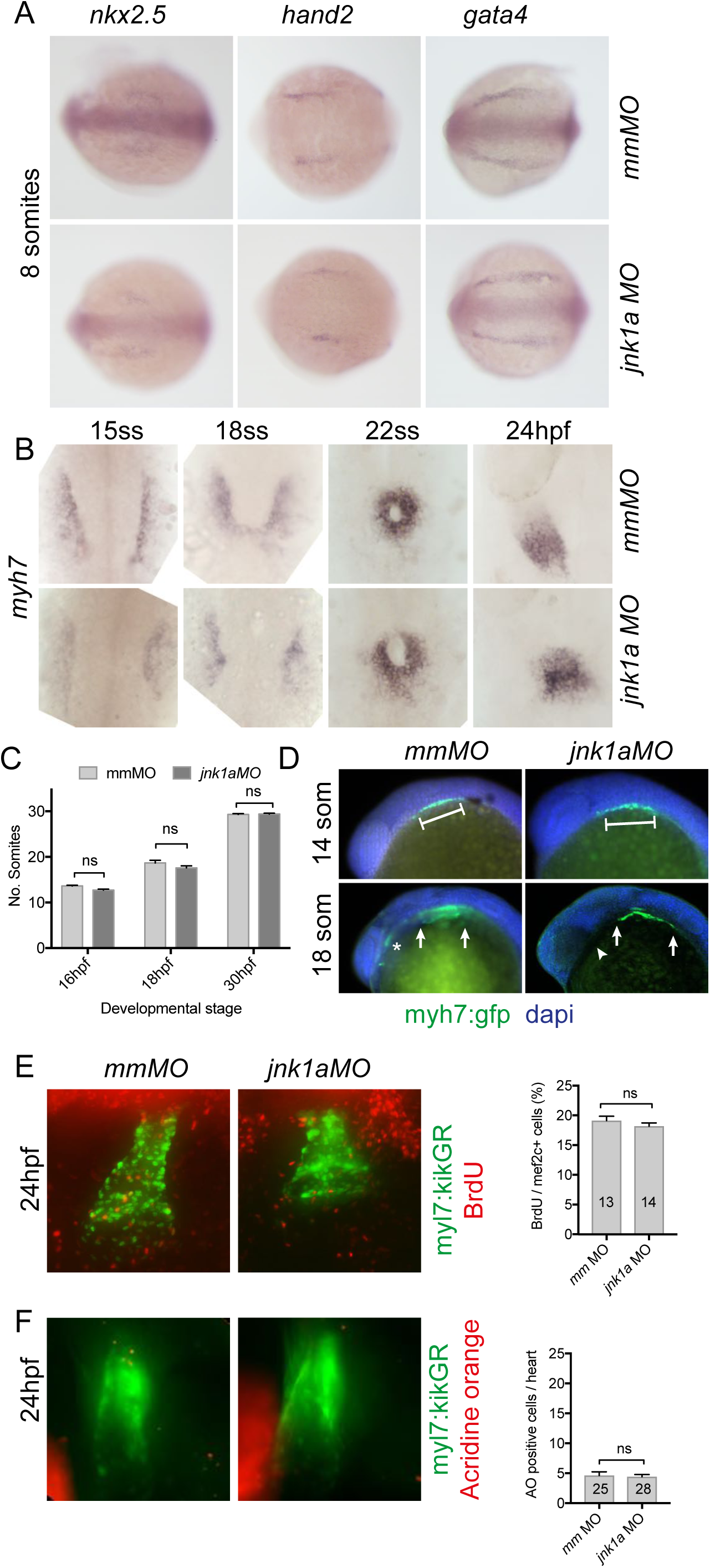
Gene expression is normal in the anterior lateral plate mesoderm. (**A**) WISH showing normal expression patterns of transcription factors required for cardiomyocyte specification in embryos injected with 2ng mmMO and 2ng *jnk1a* MO (**B**) Ventricular cardiomyocytes migrating through the anterior lateral plate mesoderm (ALPM) to form the heart are visualised by WISH for *myh7*. Although the distribution of cells is similar at 15ss between mmMO and *jnk1a* MO, there is delayed convergence at the midline in the *jnk1a* MO-treated embryos at 18 and 22ss and diminished area of *myh7* expression pattern at 24 hpf. (**C**) Lateral views of *myh7:gfp* transgenic embryos showing normal appearance of cardiomyocytes in ALPM at 14ss in *jnk1a* MO-treated embryos, but delayed migration to form the cardiac cone (*) at 18ss. (**D**) Somite counting indicates there is no developmental delay between embryos injected with 2ng mmMO and those injected with 2ng *jnk1a* MO. Proliferation (**D**) and cell death (**E**) of cardiomyocytes during migration through ALPM is not reduced in 2ng *jnk1a* morphants. ns = not significant.

Subsequently, at 22ss, there was incomplete formation of the cardiac cone in the morphants, and at 24 hpf the heart tube appeared poorly extended (Figure 5B). To exclude developmental delay as a cause of the failure to form the heart cone in the *jnk1a* morphants, we immunolabelled the forming trunk myotomes with MF20 antibody between 16 and 30 hpf. This demonstrated no difference in the number of myotomes in *jnk1a* morphants when compared with control embryos when assessed at 14ss (13 hpf), 18ss (17 hpf) or at 30hpf. (Figure 5C). The MF20 antibody also labelled cardiomyocytes and demonstrated a failure of cardiomyocytes to migrate from the lateral plate mesoderm to midline in *jnk1a* morphants. (Figure 5D). During their migration from the ALPM to the midline, cardiomyocytes undergo cell division. To identify cell that are proliferating during this phase we carried out BrdU pulsing at 16ss in *myl7:kikGR* embryos treated with *jnk1a* morpholino or mismatch control morpholino and assayed for incorporation at 28hpf. There was no difference in proliferative activity between cardiomyocytes from *jnk1a* morphants and controls (Figure 5E). Similarly, using *myl7:kikGR* in combination with acridine orange, there was no evidence of increased programmed cell or necrotic death within the heart between the morphants and the control embryos (Figure 5F).

To evaluate this abnormal cardiomyocyte migration behaviour in more detail, we carried out time-lapse confocal imaging of cardiomyocytes labelled with *myl7:gfp* between the 16ss and 20ss as they migrated from the ALPM to the midline and coalesce to form the heart cone. Close examination revealed that the migration of cardiomyocytes to the heart cone was disturbed. Tracking individual CMCs indicated the mean velocity of cells as they migrated to the heart cone was reduced in *jnk1a* morphants (Figure 6D) and this was because the cardiomyocytes took a more wandering path (Figure 6E). These abnormal behaviours were improved by co-injection of *jnk1a Ex7 Lg* mRNA, increasing their velocity (Figure 6D) and normalising the directional component of migration (Figure 6E).

**Figure 6.**
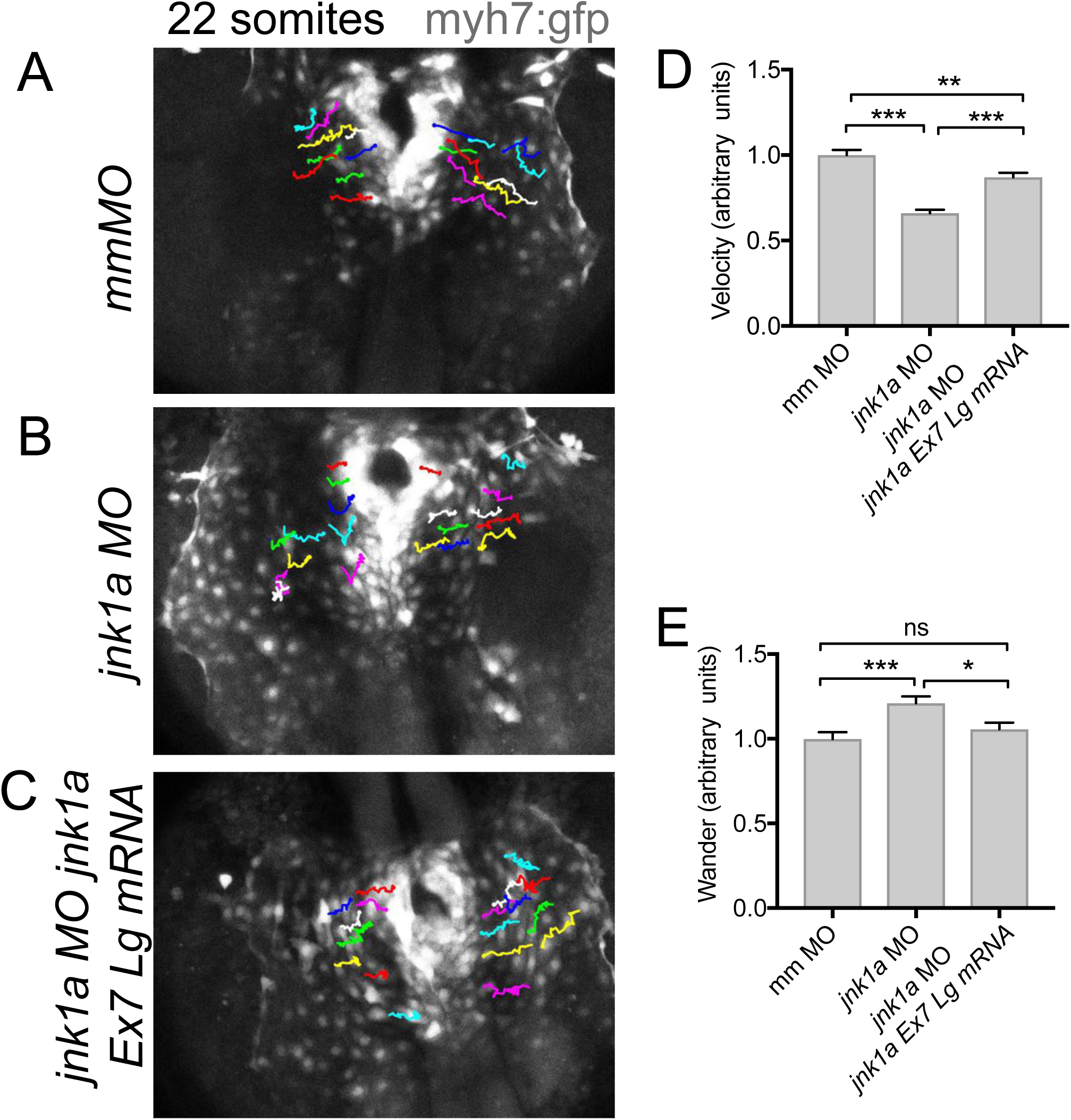
Tracking of migrating cardiomyocytes. (**A-C**) Maximal intensity projection confocal images of *myh7:gfp* labelled cardiomyocytes at 22ss in embryos injected with (**A**) 2ng mmMO (**B**) 2ng *jnk1a* MO and (**C**) 2ng *jnk1a MO* and 100pg *jnk1a Ex7 Lg* mRNA. Tracks show paths of individual cardiomyocytes and each colour represents a different track. (**D**) Mean speed and (**E**) wander index (mean speed/mean velocity) of each cardiomyocyte relative to mmMO control in embryos injected with 2ng *jnk1a* MO and also co-injected with *jnk1aEx7 Lg* mRNA. n=3 * p<0.05, ** p<0.02, ***p<0.001, ns = not significant.

## Discussion

Our understanding of heart morphogenesis and the developmental basis of congenital heart malformations is now underpinned by the concept of the first and second heart fields. The addition of SHF progenitors to the FHF-derived heart tube provides cardiac structures required for breathing air (i.e. the septated heart with a pulmonary circulation), but the presence of the SHF in zebrafish is proof that the SHF predates the evolutionary emergence of these structures [23,24,30,33].

In this study we have found that a reduction in the FHF cardiomyocyte contribution to the zebrafish ventricle occurs when *jnk1a* gene expression is abrogated and that the reduction is completely and specifically rescued by the highly conserved *jnk1a Ex7 Lg* alternative spliced transcript. Whilst the same findings in both *jnk1a* morphants and MZ null mutants, together with the highly specific mRNA rescue are compelling, suggesting that *Jnk1a* plays a fundamental role in FHF cardiogenesis, *Jnk1* null mouse embryos are not reported to have a cardiac phenotype [18,19]. However, close examination of published images indicates that *Jnk1/Jnk2* null (double mutant) mouse embryos harvested at E11.5 appear to have ceased development at E9.5, and to have a hypoplastic ventricle [19]. Detailed re-analysis of the *Jnk1*, and ideally *Jnk1/Jnk2* double null mouse embryos will be important to clarify this. Despite the lack of compensation from SHF cardiomyocytes, the reduction in the FHF-derived ventricular component is well tolerated by the *MZjnk1a* mutants. This is probably because the FHF and SHF addition add to the same chamber in fish. Notably, however, the deficiency can still be observed at 72 hpf by akinesia of the FHF-derived segment of the ventricle and reduced heart rate, which has been shown to be due to disturbed coupling between the heart field components [24]. In vertebrates with a septated heart, this 25% reduction in cardiomyocytes would be expected to be clinically apparent, as a hypoplastic left ventricle.

Despite the simplicity of the developing zebrafish heart that permits counting of individual cardiomyocytes, it is not easy to determine when or why a small group of FHF ventricular cardiomyocytes might be lost. Expression patterns of transcriptional regulators in the ALPM mark both ventricular and atrial cardiomyocytes, but also endothelial and haemopoietic lineages. Moreover, there is a small degree of cardiomyocyte proliferation during migration through the ALPM to the heart [31]. It is perhaps therefore not surprising that our WISH and proliferation assays are not sensitive enough to detect a reduction that affects less than 12 cells. Similarly, the death of a handful of cells is also difficult to determine over a window of migration from 16 hpf to 28 hpf. Importantly, we have shown that the migratory pattern of cardiomyocytes is disturbed in the absence of Jnk1a. A role for Jnk1 in cell movement is widely recognised in the metastatic spread of cancer [34] and is required in Schwann cell migration [35], dependent on paxillin at focal adhesions [36]. It is possible to imagine that some cells may become lost during migration, and in isolation undergo cell death or differentiate into other tissues. Other studies demonstrating perturbance in FHF/SHF complements [24, 37] have similarly not identified where or how cells are lost.

It has been suggested that in certain cell types, Jnk may activate AP1 and related factors [38]. In zebrafish it has been shown that AP1 disturbance affects only the SHF ventricular cardiomyocyte addition [37]. Hence *jnk1a* is unlikely to be acting through AP1 in this FHF context. Hand2 is known to control numbers of cardiomyocytes in both the FHF and SHF [39] and it is possible that *jnk1a* may be acting to augment the transcription or activity of this or an interacting transcriptional activator. Similarly, although *Jnk* has been implicated in the convergent extension movements regulated by the non-canonical Wnt PCP signalling pathway in Drosophila [40], Xenopus [41] and zebrafish [42], it appears that at least in zebrafish, *jnk1* may play a lesser role than *jnk2* [42] or *jnk3*, which may explain the lack of any overt PCP phenotype in our *MZjnk1a/MZjnk1b* mutants. Alternatively, there may be compensatory upregulation from other genes as our *jnk* mutants exhibit nonsense mediated decay (NSMD) [43]. Although we attempted pharmacological inhibition of NSMD with ethyl 2-{[(6,7-dimethyl-3-oxo-1,2,3,4-tetrahydro-2-quinoxalinyl)acetyl]amino}-4,5-dimethyl-3-thiophenecarboxylate [43] this agent was highly toxic and severely disturbed embryonic development when used within the first 24 hpf of development, which prevented examination for a more profound reduction in cardiomyocyte numbers. We can say, however, that there is no compensation from *jnk1b* transcripts as there was no additional reduction in cardiomyocytes in *MZjnk1a/MZjnk1b* mutants, and a greater reduction in cardiomyocytes was seen in *jnk1a* mutants (25%) than morphants (15%).

In the adult cardiovascular system, Jnk1 provides an important stress response to pressure overload [44] and both Jnk1 and Jnk2 have been implicated in the development of atheromatous plaques [45,46]. Our RT-PCR splicing analysis indicates that *jnk1b* transcripts become dominant in the maturing heart and it will be interesting to see whether these are up-regulated in myocardial injury as part of a stress response, or if *jnk1a Ex7 Lg* is upregulated as part of a regenerative response. However, no roles for *Jnk* in the developing heart have been previously reported. The highly specific FHF ventricular hypoplasia phenotype in zebrafish is the equivalent of left ventricular hypoplasia in the human heart. Clinically this is often recognised in conjunction with other abnormalities and presumed to be a secondary consequence, e.g. double outlet right ventricle (DORV), atrioventricular septal defect and total anomalous pulmonary venous connections. Isolated left ventricular hypoplasia is a key feature of hypoplastic left heart syndrome (HLHS), where the causes and developmental origins remain obscure [47]. Although abnormalities of the aortic valve have been suggested as a primary abnormality in HLHS, it is also possible that a primary failure of left ventricular growth could be responsible. Although easily confused with left ventricular hypoplasia with DORV, HLHS is a condition in which the aorta and pulmonary artery are normally connected to each ventricle of the heart [48]. Until these findings are modelled in mouse it is unclear whether *Jnk1* containing genetic pathways contain candidates for HLHS, or as with other PCP genes, double outlet right ventricle [6].

The importance of alternative splicing for regulating specific developmental processes remains underexplored. Whilst it has been shown to be important for regulating embryonic aspects of stem cell pluripotency and differentiation [49, 50] and has been shown to be common in the early mouse embryo [51] and in the zebrafish [52], there are few, if any, reports where specific splice variants have been shown to play a discrete mechanistic role during embryogenesis that is unique to that transcript and not shared with other related transcripts. There is tantalising circumstantial evidence in the literature to support the idea that individual transcripts may be specifically activated by different gene networks, for example the phosphatase DUSP8 preferentially inactivates exon 6a-containing (equivalent of Ex7 in zebrafish) JNK1 peptides [53]. In this study we have comprehensively shown developmental and organ specific splicing changes within the duplicated and highly conserved gene *jnk1*. Importantly, we have shown the balance of Ex7 and Ex8 expression in developing embryos is not random, as has been suggested previously [11]. Furthermore, we have shown that there is a functional interaction between central exon usage and C-terminal splicing, as only *jnk1a* mRNA containing both Ex7 and C-terminal Lg extensions can rescue ventricular hypoplasia. Taken together these studies show the importance of studying specific alternatively spliced transcripts in cardiac development and indicate the power of the zebrafish model to uncover specific and biomedically-relevant malformations, such as left ventricular hypoplasia. Population based genomic studies have been remarkably poor at discovering genetic disturbances that produce non-syndromic congenital heart disease. This study may suggest that the functional diversity afforded by alternative splicing and the regulatory mechanisms that may not immediately recognised as heart specific, may hold the key to understanding cardiac malformation.

## Materials and Methods

### Animals

Zebrafish were maintained in standard conditions [54] under the Animals (Scientific Procedures) Act 1986, United Kingdom, project license PPL6004548 and conformed to Directive 2010/63/EU of the European Parliament. All experiments were approved by the Newcastle University Animal Welfare and Ethical Review Board. Embryos were obtained from natural pairwise mating, reared in in E3 embryo medium at 28.5°C and staged by somite counting [55]. Embryos were treated with 1-phenyl 2-thiourea (PTU) from 24 hours post fertilisation (hpf) to suppress pigmentation as needed.

### Morpholino and mRNA injections

All injections were performed at the 1 cell stage. At 2 hpf non-fertilized or grossly abnormal embryos were discarded. Unless otherwise stated, 2ng of *jnk1a* morpholino, 1ng of *jnk1b* morpholino or 2ng of control morpholino were injected per embryo in each experiment (See Supplementary Table 1 for morpholino sequences).

### Mutants

Null mutant *jnk1a* and *jnk1b* zebrafish were made using CRISPR-Cas9-mediated mutagenesis [56] in the AB wildtype line with guide RNA identified using CRISPRscan [57]. Screening and subsequent genotyping was carried out by demonstrating disruption of a specific restriction site (*jnk1a*: EcorV and *jnk1b*: MwoI; see Supplementary Table 1). Mutations were confirmed by sequencing and outcrossing of lines.

### Cloning of alternatively spliced transcripts

Embryos at 24-72 hpf were pooled and RNA extracted using Trizol (Life Technologies) permitting generation of cDNA using Superscript III and oligo-dT primers. Alternatively spliced transcripts were identified by sub-cloning and sequencing of colonies (see Supplementary Table 1 for primer sequences). Transcripts for use in morpholino rescue experiments were produced using degenerate primers with at least 5 mismatches respect to the original cDNA (see Supplementary Table 1) and expressed using the zebrafish Gateway plasmid system [58]. RNA was produced using the SP6 Ambion mMessage mMachine kit (ThermoFisher Scientific). Unless otherwise stated 100pg of RNA were used in the rescue experiments.

### Semi quantitative RT-PCR splicing assay

Total RNA was extracted from 30-50 pooled embryos or 50 beating hearts [59] and cDNA produced as above. RT-PCR reactions for *jnk1a Ex7* or *Ex8* and *jnk1b Ex7* or *Ex8* were performed in triplicate with Go-taq G2 polymerase (Promega) using conditions: 95°C (2 min), [95°C (30 sec), 64°C (30 sec), 72°C (30 sec)] x 35 cycles, 72° (5 min). These conditions ensured products obtained were within the linear phase of PCR (Supplementary Figure 2). Primers are indicated in Supplementary Table 1. Products were visualised on a 2% agarose gel and gel densitometry carried out using Fiji [60]. Aliquots of each PCR product were purified by ethanol and sodium acetate precipitation and resuspended in 10µl of nuclease-free water prior to restriction enzyme digestion. Digestion for 1 hour at 37° was carried out with StyI-HF (*jnk1a*) and NheI-HF (*jnk1b*) and gel densitometry performed. These values were normalized against the *ef1α* value and with regard to the efficiency of the PCR reaction as established with plasmids containing full length transcripts (Supplementary Figure 2).

### Wholemount in-situ hybridisation

Full-length *jnk1a* and *jnk1b* alternatively spliced transcripts were used to generate riboprobes. For *jnk2* and *jnk3* an 1100 amplicon was selected. In addition: *hand2, nkx2.5, vmhc* (Deborah Yelon, UCSF, USA), *gata4* (Roger Patient, Oxford University, UK), *southpaw*, (Steve Wilson, UCL, UK) were generously provided by the indicated researchers. Chromogenic *in situ* hybridization was performed according to established protocols [61].

### Immunohistochemistry

Whole-mount immunohistochemistry was performed as previously described [62]. Primary antibodies: chicken anti-GFP (1/500; ab13970, Abcam), mouse anti MF20 (1/10; MYH1, DSHB), mouse anti S46 (1/10; MYHCA, DSHB), rabbit anti Mef2 (1/100; ab646444, Abcam), rat anti BrdU (1-100; ab6326, Abcam). Secondary antibodies: AlexaFluor-488 anti-mouse (1/300; A-21202, Invitrogen), AlexaFluor-488 anti-mouse IgG1 (1/300; A-21121, Invitrogen), AlexaFluor-488 anti-chicken (1/300; A11039 Invitrogen), AlexaFluor-568 anti-mouse IgG1 (1/300; A-21124, Invitrogen), AlexaFluor-568 anti-mouse IgG2b (1/300; A-21144, Invitrogen), AlexaFluor-568 anti-rabbit (1/300; A10042, Invitrogen), AlexaFluor-568 anti-rat (1/300; A11077, Invitrogen). Nuclei were counterstained with DAPI (1:10,000; D9542, Sigma).

### Cardiomyocyte quantification

Cardiomyocyte nuclei were identified by immunolabelling with Mef2 antibody at 28 hpf [33] and the *myh7:DsNucRed* transgene at 52 hpf [32]. Atrial cardiomyocytes were identified by co-labelling with S46 antibody. Positioning under a glass coverslip allowed all cardiomyocyte nuclei to be imaged in the same focal plane and manual counting was performed using Fiji [60]. To differentiate the first heart field cardiomyocyte complement from the second heart field accretion we used we used *myh7:kikGR* transgenic line injected with either mismatch control or *jnk1a* morpholino. Embryos were exposed to UV light at 20hpf, photoconverting all cardiomyocytes already expressing kikGR from green to red emitting forms. Imaging and quantification were performed as before.

### Live imaging of embryonic heart activity

Embryos were orientated in 1% low melting point agarose containing 0.2% Tricaine. Movies were taken on an inverted microscope with DIC (Nikon Diaphot) using a high-speed digital video camera (640×480 pixels at 127 frames/sec) (Multipix Imaging, Hampshire, UK). Data was analysed using Fiji [60] and M-mode obtained by reslice through the image series.

### Proliferation assay

Control or *jnk1a* morphant embryos (*myh7:gfp*) were incubated in 10mM BrdU (B5002, Sigma) in 15% DMSO for 30 min on ice at 16 somites. Then, embryos were washed in E3 medium and allowed to develop until 28 hpf. After fixing in PFA 4% overnight at 4°C, embryos were dehydrated and stored in methanol at −20°C until further use. Embryos were rehydrated in PBS, then permeabilized using Proteinase K and post-fixed in PFA 4%. After incubation in 2N HCl for 1 hour, standard IHC using anti BrdU and anti GFP antibodies was carried out. The proportion of BrdU positive cardiomyocytes was assessed as above.

### Cell death

Embryos from the *myh7:kikgr* line were injected with either control or *jnk1a* morpholino and incubated in 10µg/ml of acridine orange during the phase of migration from ALPM to the cardiac cone. As acridine orange is visualised in the FITC fluorescent channel, photoconversion of kikGR in FHF cardiomyocytes allowed identification of cardiomyocyte nuclei using the TRITC channel. As above, acridine orange positive cardiomyocytes were quantified using Fiji [60].

### Cardiomyocyte tracking

An established protocol [63] with minor modification was used to track the migration of the cardiomyocytes from the anterolateral plate mesoderm to the cardiac cone. *myh7:gfp* embryos at the 14 somite stage (ss) were hand dechorionated and oriented in 1.5% low melting agarose. A 20-image stack (A1R confocal Microscope, Nikon), using a 20x objective at 3um intervals, was acquired every 6 minutes between 15-16 ss to 19-20 ss. Manual tracking was performed using Fiji [60] and XY coordinates of individual cardiomyocytes were established for every time point. Based on these coordinates we calculated the following parameters:

Mean speed calculated from each measurable displacement per unit time between the initial (x_i_, y_i_) and the final (x_f_, y_f_) coordinates

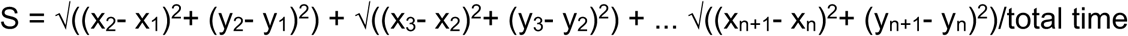

Mean velocity calculated from the absolute displacement per unit time between initial (x_i_, y_i_) and the final (x_f_, y_f_) coordinates

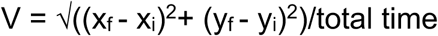

Wandering Index: Ratio between speed and velocity

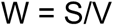

## Statistics

Where possible, sample size was determined with regard to normal values and standard deviations described in previously published experiments. In all cases at least three independent experiments, using embryos from three different clutches of eggs, were performed. Statistics were performed using Graphpad Prism7 for Mac OSX, version 6.0c (Graphpad Software Inc, USA) and, depending on the number of groups to compare, either Student t test or ANOVA was performed.

## Supporting information

control tracking

jnk1a tracking

rescue tracking

## Acknowledgments

The authors would like to thank Professor David Elliot, Newcastle University, UK for helpful advice in designing the RT-PCR splicing assay.

