## Supplementary Figures for "An alternatively spliced zebrafish *jnk1a* transcript has an essential and non-redundant role in development of the first heart field derived proximal ventricular chamber"

Suplementary figure 1

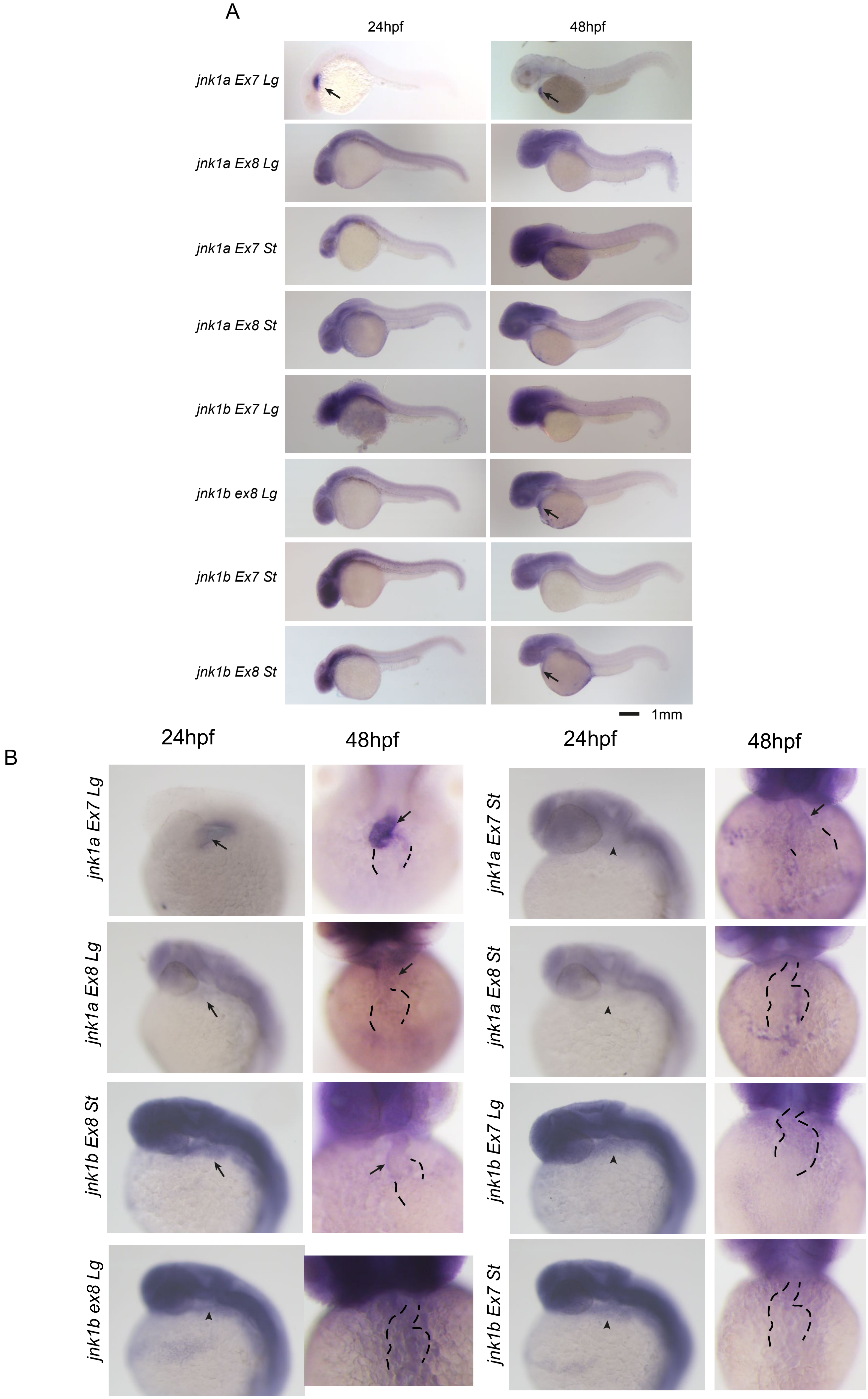

Suplementary figure 2

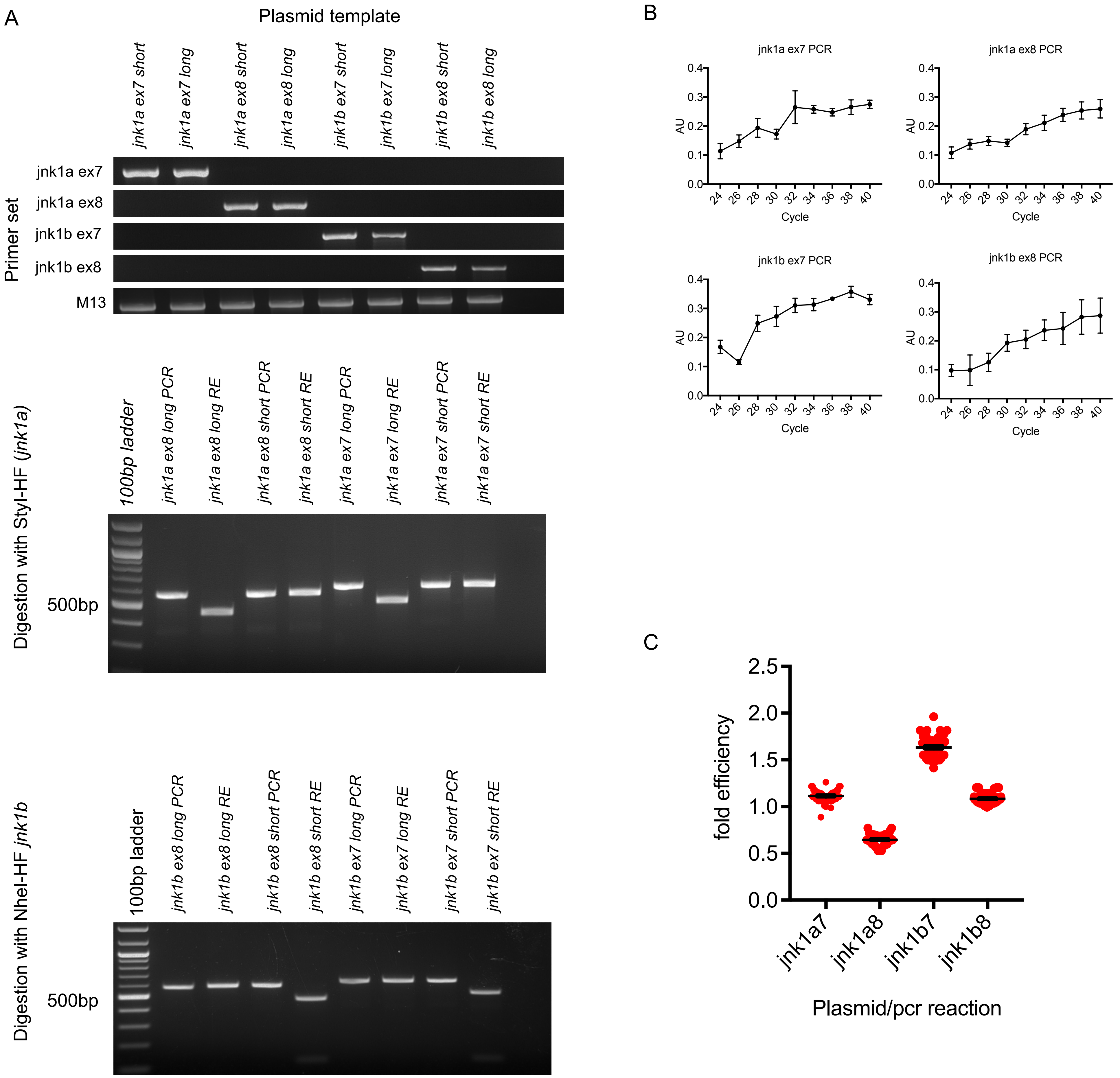

**Supplementary legends**

**Supplementary figure 1: WISH for *jnk* transcripts.** (**A**) WISH for all *jnk1a* and *jnk1b* alternatively spliced transcripts at 24 hpf and 48 hpf. Transcripts with obvious expression within heart are indicated (arrow). (**B**) Close up of heart in all *jnk1a* transcripts and 24 and 48 hpf. Expression in heart indicated by arrows and absent expression indicated by arrowheads. Outline of heart indicated by dashed lines.

**Supplementary figure 2: Validation of Splicing assay** (**A**) specificity of primers used in RT-PCR splicing assay. Primers sets to identify Ex7 and Ex 8 containing variants of both *jnk1a* and *jnk1b* transcripts were assessed against all 8 transcripts contained within plasmids, using primers for the plasmid backbone (M13) as loading control. Restriction enzyme digests showing original PCR product and complete digestion by enzyme to identify C-terminal extension in *jnk1a* and *jnk1b* transcripts. (**B**) Gel densitometry indicating splicing assay was carried out during linear phase of amplification for jnk 1a and jnk1b ex7 and ex8 PCR (Arbitrary logarithmic units). (**C**), PCR with plasmid containing transcripts were used as positive controls for PCR reactions and indicated efficiency of each reaction.

**Supplementary table1. Sequences of oligonucleotides used in study**

| **GuideRNA** | |
|---|---|
| *jnk1a gRNA* | ATTTAGGTGACACTATAGGAATGGGATATCAAGCCAAGTTTTAGAGCTAGAAATAGCAAG |
| *jnk1b gRNA* | ATTTAGGTGACACTATAGGAGGCCGGTTGCAGCCGTTGTTTTAGAGCTAGAAATAGCAAG |

| **Genotyping primers** | |
|---|---|
| *jnk1a$^{n1}$ fwd* | GCTACAGGTCTGCTGATGACAC |
| *jnk1a$^{n1}$ rev* | TTTTGATGCTGAAACCACAAAG |
| *jnk1b$^{n2}$ fwd* | TTTTCCTAGGATCTGAAACCA |
| *jnk1b$^{n2}$ rev* | CCGTTTGGTGAGATCTGTTTTC |

| **Transcript cloning** | |
|---|---|
| *jnk1a ATG fwd* | ATGAACAAAAATAAGCGAGA |
| *jnk1a STOP long rev* | TCATCTGCAGCAGCTCAGGG |
| *jnk1b ATG fwd* | ATGAACAGGAATAAGCGCGA |
| *jnk1b STOP long rev* | TCATCTGCAGCAGTGCAGCG |
| *jnk1a and b STOP short common rev* | TCACTGCTGCACCTGTGCTA |

| | **Morpholino resistant rescue transcripts** |
|---|---|
| *jnk1a rescue fwd* | GGGGACAAGTTTGTACAAAAAAGCAGGCTACATGAATAAGAACAAGAGGGAGAAGGAATTC |
| *jnk1a rescue long rev* | GGGGACCACTTTGTACAAGAAAGCTGGGTGTCATCTGCAGCAGCTCAGGGGTCCCG |
| *jnk1b rescue fwd* | GGGGACAAGTTTGTACAAAAAAGCAGGCTACATGAATCGCAACAAACGGGAGAAGGAAT |
| *jnk1b rescue long rev* | GGGGACCACTTTGTACAAGAAAGCTGGGTGTCATCTGCAGCAGTGCAGCGCTCCG |
| *jnk1a and b rescue short rev common* | GGGGACCACTTTGTACAAGAAAGCTGGGTGTCACTGCTGCACCTGTGCTA |

| | **RT-PCR splicing assay** |
|---|---|
| *jnk1a ex7 fwd* | TGTGGACATTTGGTCTGTGG |

| | |
|---|---|
| *jnk1a ex8 fwd* | TTTCCGGGTTCAGATCATATTGA |
| *jnk1a common rev* | AAGCTGCTGTCTGTGTCTGA |
| *jnk1b ex7 fwd* | ACGTGGATATTTGGGCTGTTG |
| *jnk1b ex8 fwd* | AGTGTGTTGTTTCCTGGCAC |
| *jnk1b common rev* | ACTGCTGTCGGTGTCTGAG |
| *ef1αforw* | CTTCTCAGGCTGACTGTGC |
| *ef1αrev* | CCGCTAGCATTACCCTCC |

| | **RT-PCR** |
|---|---|
| *jnk2 fwd* | CCACCTTTACCGTCCTCAGA |
| *jnk2 rev* | CAATGCAAGAGCTGTCGGTA |
| *jnk3 fwd* | ACCTTCACGGTTCTCAAACG |
| *jnk3 rev* | CCCTTCACCACACCATTCTT |

| | ***In situ* hybridization** |
|---|---|
| *jnk2 fwd-T7* | taatacgactcactatagggCTCCACCTTTACCGTCCTCA |
| *jnk2 rev-T3* | attaaccctcactaaagggaCGTCGACATGGATGAGATGT |
| *jnk3 fwd-T7* | taatacgactcactatagggTGAGCAAAAGCAAAGTGGAC |
| *jnk3 rev-T3* | attaaccctcactaaagggaCTCTCCTCGAAGTTCATCACC |

| | **Morpholino Sequences** |
|---|---|
| *jnk1a MO* | CTCTCGCTTATTTTTGTTCATGGTG |
| *jnk1b MO* | CTTTCTCGCGCTTATTCCTGTTCAT |
| *control MO* | CCTCTTACCTCAGTTACAATTTATA |

**Supplementary table 2**

**Sequences of jnk1a and jnk1b alternatively spliced transcripts.**

>jnk1a_ex7_sh
ATGAACAAAAATAAGCGAGAGAAAGAATTCTACAGTGTAGATGTGGGGGATTCGACA
TTCACAGTGTTGAAGCGCTATCAGAATCTAAGACCCATTGGCTCGGGTGCTCAGGGA
ATAGTCTGCTCAGCGTATGACAACAACCTCGAGCGAAACGTGGCTATAAAGAAACTC
AGCCGGCCTTTTCAAAATCAAACTCATGCCAAACGTGCGTACAGAGAGCTGGTGCTC
ATGAAATGTGTCAACCATAAAAATATAATAGGCCTCTTAAATGTTTTTACACCGCAA
AAAACATTAGAAGAATTCCAAGATGTTTACCTAGTAATGGAGCTGATGGATGCGAAC
CTCTGCCAAGTCATTCAGATGGAGCTGGACCACGAGCGGCTGTCCTATCTGCTGTAC
CAGATGCTGTGTGGAATAAAACACCTCCACGCGGCGGGGATCATCCACAGGGACCTG
AAACCCAGTAACATCGTGGTAAAGTCAGACTGTACCCTGAAGATCCTGGATTTCGGT
CTGGCGCGGACAGCAGCTACAGGTCTGCTGATGACACCATATGTGGTGACCCGTTAC
TACAGAGCCCCTGAAGTCATCCTGGGAATGGGATATCAAGCCAATGTGGACATTTGG
TCTGTGGGCTGCATTTTGGCAGAAATGGTCCGTCACAAAATCCTTTTTCCTGGGAGG
GACTATATTGATCAGTGGAATAAAGTAATAGAGCAGCTGGGAACGCCAACTCAGGAG
TTCTTGTTGAAACTCAACCAGTCTGTGCGGACCTATGTGGAGAACAGGCCCCGGTAC
ACTGGATATAGCTTTGAGAAGCTGTTTCCTGATGTCCTGTTCCCTGCTGATTCAGAA
CACAGCAAACTAAAAGCGAGTCAGGCGCGGGACCTGCTGTCTAAAATGCTGGTGATT
GATGCATCAAAACGAATCTCGGTGGATGAGGCTTTGCAGCACCCCTACATTAACGTG
TGGTACGACCCGGCTGAAGTGGAAGCGCCTTCTCCTCTGATCACAGACAAACAGCTC
GATGAGAGGGAACACACAGTGGAAGAGTGGAAAGAACTGATCTATAAAGAAGTGCTG
GATTGGGAAGAACGGATGAAGAACGGTGTTATTCGAGGTCAGCCCTCCCCCCTAGCA
CAGGTGCAGCAGTGA

> jnk1a_ex7_sh Protein
MNKNKREKEFYSVDVGDSTFTVLKRYQNLRPIGSGAQGIVCSAYDNNLERNVAIKKL
SRPFQNQTHAKRAYRELVLMKCVNHKNIIGLLNVFTPQKTLEEFQDVYLVMELMDAN
LCQVIQMELDHERLSYLLYQMLCGIKHLHAAGIIHRDLKPSNIVVKSDCTLKILDFG
LARTAATGLLMTPYVVTRYYRAPEVILGMGYQANVDIWSVGCILAEMVRHKILFPGR
DYIDQWNKVIEQLGTPTQEFLLKLNQSVRTYVENRPRYTGYSFEKLFPDVLFPADSE
HSKLKASQARDLLSKMLVIDASKRISVDEALQHPYINVWYDPAEVEAPSPLITDKQL
DEREHTVEEWKELIYKEVLDWEERMKNGVIRGQPSPLAQVQQ*

```
>jnk1a_ex7_Lg
ATGAACAAAAATAAGCGAGAGAAAGAATTCTACAGTGTAGATGTGGGGGATTCGACA
TTCACAGTGTTGAAGCGCTATCAGAATCTAAGACCCATTGGCTCGGGTGCTCAGGGA
ATAGTCTGCTCAGCGTATGACAACAACCTCGAGCGAAACGTGGCTATAAAGAAACTC
AGCCGGCCTTTTCAAAATCAAACTCATGCCAAACGTGCGTACAGAGAGCTGGTGCTC
ATGAAATGTGTCAACCATAAAAATATAATAGGCCTCTTAAATGTTTTTACACCGCAA
AAAACATTAGAAGAATTCCAAGATGTTTACCTAGTAATGGAGCTGATGGATGCGAAC
CTCTGCCAAGTCATTCAGATGGAGCTGGACCACGAGCGGCTGTCCTATCTGCTGTAC
CAGATGCTGTGTGGAATAAAACACCTCCACGCGGCGGGGATCATCCACAGGGACCTG
AAACCCAGTAACATCGTGGTAAAGTCAGACTGTACCCTGAAGATCCTGGATTTCGGT
CTGGCGCGGACAGCAGCTACAGGTCTGCTGATGACACCATATGTGGTGACCCGTTAC
TACAGAGCCCCTGAAGTCATCCTGGGAATGGGATATCAAGCCAATGTGGACATTTGG
TCTGTGGGCTGCATTTTGGCAGAAATGGTCCGTCACAAAATCCTTTTTCCTGGGAGG
GACTATATTGATCAGTGGAATAAAGTAATAGAGCAGCTGGGAACGCCAACTCAGGAG
TTCTTGTTGAAACTCAACCAGTCTGTGCGGACCTATGTGGAGAACAGGCCCCGGTAC
ACTGGATATAGCTTTGAGAAGCTGTTTCCTGATGTCCTGTTCCCTGCTGATTCAGAA
CACAGCAAACTAAAAGCGAGTCAGGCGCGGGACCTGCTGTCTAAAATGCTGGTGATT
GATGCATCAAAACGAATCTCGGTGGATGAGGCTTTGCAGCACCCCTACATTAACGTG
TGGTACGACCCGGCTGAAGTGGAAGCGCCTTCTCCTCTGATCACAGACAAACAGCTC
GATGAGAGGGAACACACAGTGGAAGAGTGGAAAGAACTGATCTATAAAGAAGTGCTG
GATTGGGAAGAACGGATGAAGAACGGTGTTATTCGAGGTCAGCCCTCCCCCCTAGGT
GCAGCAGTGATCAACGGCTCACCCCAGCCCTCATCCTCATCCTCCATCAACGACGTG
TCCTCCATGTCCACAGAGCCCACCGTGGCCTCAGACACAGACAGCAGCTTAGAGGCC
TCGGCGGGACCCCTGAGCTGCTGCAGATGA

> jnk1a_ex7_Lg Protein
MNKNKREKEFYSVDVGDSTFTVLKRYQNLRPIGSGAQGIVCSAYDNNLERNVAIKKL
SRPFQNQTHAKRAYRELVLMKCVNHKNIIGLLNVFTPQKTLEEFQDVYLVMELMDAN
LCQVIQMELDHERLSYLLYQMLCGIKHLHAAGIIHRDLKPSNIVVKSDCTLKILDFG
LARTAATGLLMTPYVVTRYYRAPEVILGMGYQANVDIWSVGCILAEMVRHKILFPGR
DYIDQWNKVIEQLGTPTQEFLLKLNQSVRTYVENRPRYTGYSFEKLFPDVLFPADSE
HSKLKASQARDLLSKMLVIDASKRISVDEALQHPYINVWYDPAEVEAPSPLITDKQL
DEREHTVEEWKELIYKEVLDWEERMKNGVIRGQPSPLGAAVINGSPQPSSSSSINDV
SSMSTEPTVASDTDSSLEASAGPLSCCR*
```

```
>jnk1a_ex8_sh
ATGAACAAAAATAAGCGAGAGAAAGAATTCTACAGTGTAGATGTGGGGGATTCGACA
TTCACAGTGTTGAAGCGCTATCAGAATCTAAGACCCATTGGCTCGGGTGCTCAGGGA
ATAGTCTGCTCAGCGTATGACAACAACCTCGAGCGAAACGTGGCTATAAAGAAACTC
AGCCGGCCTTTTCAAAATCAAACTCATGCCAAACGTGCGTACAGAGAGCTGGTGCTC
ATGAAATGTGTCAACCATAAAAATATAATAGGCCTCTTAAATGTTTTTACACCGCAA
AAAACATTAGAAGAATTCCAAGATGTTTACCTAGTAATGGAGCTGATGGATGCGAAC
CTCTGCCAAGTCATTCAGATGGAGCTGGACCACGAGCGGCTGTCCTATCTGCTGTAC
CAGATGCTGTGTGGAATAAAACACCTCCACGCGGCGGGGATCATCCACAGGGACCTG
AAACCCAGTAACATCGTGGTAAAGTCAGACTGTACCCTGAAGATCCTGGATTTCGGT
CTGGCGCGGACAGCAGCTACAGGTCTGCTGATGACACCATATGTGGTGACCCGTTAC
TACAGAGCCCCTGAAGTCATCCTGGGAATGGGATATCAAGCCAATGTGGATGTGTGG
TCTGTCGGCTGTATCATGGCTGAAATGGTCAGAGGTAGTGTATTATTTCCGGGTTCA
GATCATATTGATCAGTGGAATAAAGTAATAGAGCAGCTGGGAACGCCAACTCAGGAG
TTCTTGTTGAAACTCAACCAGTCTGTGCGGACCTATGTGGAGAACAGGCCCCGGTAC
ACTGGATATAGCTTTGAGAAGCTGTTTCCTGATGTCCTGTTCCCTGCTGATTCAGAA
CACAGCAAACTAAAAGCGAGTCAGGCGCGGGACCTGCTGTCTAAAATGCTGGTGATT
GATGCATCAAAACGAATCTCGGTGGATGAGGCTTTGCAGCACCCCTACATTAACGTG
TGGTACGACCCGGCTGAAGTGGAAGCGCCTTCTCCTCTGATCACAGACAAACAGCTC
GATGAGAGGGAACACACAGTGGAAGAGTGGAAAGAACTGATCTATAAAGAAGTGCTG
GATTGGGAAGAACGGATGAAGAACGGTGTTATTCGAGGTCAGCCCTCCCCCCTAGCA
CAGGTGCAGCAGTGA

> jnk1a_ex8_sh protein
MNKNKREKEFYSVDVGDSTFTVLKRYQNLRPIGSGAQGIVCSAYDNNLERNVAIKKL
SRPFQNQTHAKRAYRELVLMKCVNHKNIIGLLNVFTPQKTLEEFQDVYLVMELMDAN
LCQVIQMELDHERLSYLLYQMLCGIKHLHAAGIIHRDLKPSNIVVKSDCTLKILDFG
LARTAATGLLMTPYVVTRYYRAPEVILGMGYQANVDVWSVGCIMAEMVRGSVLFPGS
DHIDQWNKVIEQLGTPTQEFLLKLNQSVRTYVENRPRYTGYSFEKLFPDVLFPADSE
HSKLKASQARDLLSKMLVIDASKRISVDEALQHPYINVWYDPAEVEAPSPLITDKQL
DEREHTVEEWKELIYKEVLDWEE
```

```
>jnk1a_ex8_Lg
ATGAACAAAAATAAGCGAGAGAAAGAATTCTACAGTGTAGATGTGGGGGATTCGACA
TTCACAGTGTTGAAGCGCTATCAGAATCTAAGACCCATTGGCTCGGGTGCTCAGGGA
ATAGTCTGCTCAGCGTATGACAACAACCTCGAGCGAAACGTGGCTATAAAGAAACTC
AGCCGGCCTTTTCAAAATCAAACTCATGCCAAACGTGCGTACAGAGAGCTGGTGCTC
ATGAAATGTGTCAACCATAAAAATATAATAGGCCTCTTAAATGTTTTTACACCGCAA
AAAACATTAGAAGAATTCCAAGATGTTTACCTAGTAATGGAGCTGATGGATGCGAAC
CTCTGCCAAGTCATTCAGATGGAGCTGGACCACGAGCGGCTGTCCTATCTGCTGTAC
CAGATGCTGTGTGGAATAAAACACCTCCACGCGGCGGGGATCATCCACAGGGACCTG
AAACCCAGTAACATCGTGGTAAAGTCAGACTGTACCCTGAAGATCCTGGATTTCGGT
CTGGCGCGGACAGCAGCTACAGGTCTGCTGATGACACCATATGTGGTGACCCGTTAC
TACAGAGCCCCTGAAGTCATCCTGGGAATGGGATATCAAGCCAATGTGGATGTGTGG
TCTGTCGGCTGTATCATGGCTGAAATGGTCAGAGGTAGTGTATTATTTCCGGGTTCA
GATCATATTGATCAGTGGAATAAAGTAATAGAGCAGCTGGGAACGCCAACTCAGGAG
TTCTTGTTGAAACTCAACCAGTCTGTGCGGACCTATGTGGAGAACAGGCCCCGGTAC
ACTGGATATAGCTTTGAGAAGCTGTTTCCTGATGTCCTGTTCCCTGCTGATTCAGAA
CACAGCAAACTAAAAGCGAGTCAGGCGCGGGACCTGCTGTCTAAAATGCTGGTGATT
GATGCATCAAAACGAATCTCGGTGGATGAGGCTTTGCAGCACCCCTACATTAACGTG
TGGTACGACCCGGCTGAAGTGGAAGCGCCTTCTCCTCTGATCACAGACAAACAGCTC
GATGAGAGGGAACACACAGTGGAAGAGTGGAAAGAACTGATCTATAAAGAAGTGCTG
GATTGGGAAGAACGGATGAAGAACGGTGTTATTCGAGGTCAGCCCTCCCCCCTAGGT
GCAGCAGTGATCAACGGCTCACCCCAGCCCTCATCCTCATCCTCCATCAACGACGTG
TCCTCCATGTCCACAGAGCCCACCGTGGCCTCAGACACAGACAGCAGCTTAGAGGCC
TCGGCGGGACCCCTGAGCTGCTGCAGATGA

> jnk1a_ex8_Lg Protein
MNKNKREKEFYSVDVGDSTFTVLKRYQNLRPIGSGAQGIVCSAYDNNLERNVAIKKL
SRPFQNQTHAKRAYRELVLMKCVNHKNIIGLLNVFTPQKTLEEFQDVYLVMELMDAN
LCQVIQMELDHERLSYLLYQMLCGIKHLHAAGIIHRDLKPSNIVVKSDCTLKILDFG
LARTAATGLLMTPYVVTRYYRAPEVILGMGYQANVDVWSVGCIMAEMVRGSVLFPGS
DHIDQWNKVIEQLGTPTQEFLLKLNQSVRTYVENRPRYTGYSFEKLFPDVLFPADSE
HSKLKASQARDLLSKMLVIDASKRISVDEALQHPYINVWYDPAEVEAPSPLITDKQL
DEREHTVEEWKELIYKEVLDWEERMKNGVIRGQPSPLGAAVINGSPQPSSSSSINDV
SSMSTEPTVASDTDSSLEASAGPLSCCR*
```

```
>jnk1b_ex7_sh
ATGAACAGGAATAAGCGCGAGAAAGAATATTACAGCATAGATGTAGGAGATTCGACG
TTCACCGTTTTGAAGCGCTATCAGAATTTAAGACCAATCGGGTCCGGAGCACAAGGC
ATCGTCTGCTCAGCGTATGACCACGTCCTCGATCGAAATGTGGCGATTAAGAAACTC
AGCCGACCCTTTCAAAACCAAACTCATGCCAAACGGGCCTACAGAGAACTGGTCCTG
ATGAAATGCGTCAACCACAAAAATATAATTGGCTTACTAAACGTGTTCACACCACAG
AAGACCCTTGAAGAGTTCCAGGATGTTTATCTGGTGATGGAGCTGATGGATGCAAAC
CTGTGTCAGGTGATTCAGATGGAGCTGGACCACGAGAGGCTGTCCTACCTGCTCTAT
CAGATGCTCTGCGGCATTAAACACCTGCACGCTGCTGGCATCATACACAGGGACCTG
AAACCCAGTAATATAGTAGTGAAATCGGACTGCACGCTGAAGATCCTGGATTTCGGT
CTGGCCAGAACGGCTGCAACCGGCCTCCTCATGACTCCTTATGTAGTGACACGCTAT
TATCGGGCCCCAGAGGTCATCCTGGGCATGGGTTATCAAGCTAACGTGGATATTTGG
GCTGTTGGCTGCATTATGGCAGAGATGGTGCGGCACAAAATCCTTTTTCCAGGGAGG
GACTATATTGACCAGTGGAATAAAGTGATCGAGCAGCTCGGCACGCCGTCACAGGAG
TTCATGATGAAGCTGAATCAGTCTGTGAGGACGTATGTGGAGAACCGGCCTCGGTAT
GCGGGATACAGCTTTGAGAAGCTCTTCCCAGACGTGCTCTTCCCCGCAGACTCGGAC
CACAACAAACTCAAGGCGAGTCAGGCACGAGACTTGTTATCCAAAATGCTGGTAATA
GATGCGTCCAAGCGGATCTCTGTAGACGAGGCGCTTCAGCACCCCTACATCAACGTT
TGGTACGACCCGTCAGAAGTGGAGGCGCCACCACCAGCGATCACGGATAAACAGCTC
GATGAGAGAGAACACTCAGTGGAAGAGTGGAAAGAGCTCATATATAAGGAAGTGCTG
GAATGGGAGGAGCGAACAAAAAATGGAGTGATCAGAGGACAGCCGGCCTCGCTAGCA
CAGGTGCAGCAGTGA

> jnk1b_ex7_sh Protein
MNRNKREKEYYSIDVGDSTFTVLKRYQNLRPIGSGAQGIVCSAYDHVLDRNVAIKKL
SRPFQNQTHAKRAYRELVLMKCVNHKNIIGLLNVFTPQKTLEEFQDVYLVMELMDAN
LCQVIQMELDHERLSYLLYQMLCGIKHLHAAGIIHRDLKPSNIVVKSDCTLKILDFG
LARTAATGLLMTPYVVTRYYRAPEVILGMGYQANVDIWAVGCIMAEMVRHKILFPGR
DYIDQWNKVIEQLGTPSQEFMMKLNQSVRTYVENRPRYAGYSFEKLFPDVLFPADSD
HNKLKASQARDLLSKMLVIDASKRISVDEALQHPYINVWYDPSEVEAPPPAITDKQL
DEREHSVEEWKELIYKEVLEWEERTKNGVIRGQPASLAQVQQ*
```

>jnk1b_ex7_Lg
ATGAACAGGAATAAGCGCGAGAAAGAATATTACAGCATAGATGTAGGAGATTCGACG
TTCACCGTTTTGAAGCGCTATCAGAATTTAAGACCAATCGGGTCCGGAGCACAAGGC
ATCGTCTGCTCAGCGTATGACCACGTCCTCGATCGAAATGTGGCGATTAAGAAACTC
AGCCGACCCTTTCAAAACCAAACTCATGCCAAACGGGCCTACAGAGAACTGGTCCTG
ATGAAATGCGTCAACCACAAAAATATAATTGGCTTACTAAACGTGTTCACACCACAG
AAGACCCTTGAAGAGTTCCAGGATGTTTATCTGGTGATGGAGCTGATGGATGCAAAC
CTGTGTCAGGTGATTCAGATGGAGCTGGACCACGAGAGGCTGTCCTACCTGCTCTAT
CAGATGCTCTGCGGCATTAAACACCTGCACGCTGCTGGCATCATACACAGGGACCTG
AAACCCAGTAATATAGTAGTGAAATCGGACTGCACGCTGAAGATCCTGGATTTCGGT
CTGGCCAGAACGGCTGCAACCGGCCTCCTCATGACTCCTTATGTAGTGACACGCTAT
TATCGGGCCCCAGAGGTCATCCTGGGCATGGGTTATCAAGCTAACGTGGATATTTGG
GCTGTTGGCTGCATTATGGCAGAGATGGTGCGGCACAAAATCCTTTTTCCAGGGAGG
GACTATATTGACCAGTGGAATAAAGTGATCGAGCAGCTCGGCACGCCGTCACAGGAG
TTCATGATGAAGCTGAATCAGTCTGTGAGGACGTATGTGGAGAACCGGCCTCGGTAT
GCGGGATACAGCTTTGAGAAGCTCTTCCCAGACGTGCTCTTCCCCGCAGACTCGGAC
CACAACAAACTCAAGGCGAGTCAGGCACGAGACTTGTTATCCAAAATGCTGGTAATA
GATGCGTCCAAGCGGATCTCTGTAGACGAGGCGCTTCAGCACCCCTACATCAACGTT
TGGTACGACCCGTCAGAAGTGGAGGCGCCACCACCAGCGATCACGGATAAACAGCTC
GATGAGAGAGAACACTCAGTGGAAGAGTGGAAAGAGCTCATATATAAGGAAGTGCTG
GAATGGGAGGAGCGAACAAAAAATGGAGTGATCAGAGGACAGCCGGCCTCGCTAGGT
GCAGCAGTGAGCAGTGACTCCCATGAGCCCTCGACGTCGTCCTCCTCCATAAACGAT
GTGTCGTCCATGTCCACCGAGGTCACGCTGACCTCAGACACCGACAGCAGTCAGGAG
ACGTCCAACGGAGCGCTGCACTGCTGCAGATGA

> jnk1b_ex7_Lg Protein
MNRNKREKEYYSIDVGDSTFTVLKRYQNLRPIGSGAQGIVCSAYDHVLDRNVAIKKL
SRPFQNQTHAKRAYRELVLMKCVNHKNIIGLLNVFTPQKTLEEFQDVYLVMELMDAN
LCQVIQMELDHERLSYLLYQMLCGIKHLHAAGIIHRDLKPSNIVVKSDCTLKILDFG
LARTAATGLLMTPYVVTRYYRAPEVILGMGYQANVDIWAVGCIMAEMVRHKILFPGR
DYIDQWNKVIEQLGTPSQEFMMKLNQSVRTYVENRPRYAGYSFEKLFPDVLFPADSD
HNKLKASQARDLLSKMLVIDASKRISVDEALQHPYINVWYDPSEVEAPPPAITDKQL
DEREHSVEEWKELIYKEVLEWEERTKNGVIRGQPASLGAAVSSDSHEPSTSSSSIND
VSSMSTEVTLTSDTDSSQETSNGALHCCR*

```
>jnk1b_ex8_sh
ATGAACAGGAATAAGCGCGAGAAAGAATATTACAGCATAGATGTAGGAGATTCGACG
TTCACCGTTTTGAAGCGCTATCAGAATTTAAGACCAATCGGGTCCGGAGCACAAGGC
ATCGTCTGCTCAGCGTATGACCACGTCCTCGATCGAAATGTGGCGATTAAGAAACTC
AGCCGACCCTTTCAAAACCAAACTCATGCCAAACGGGCCTACAGAGAACTGGTCCTG
ATGAAATGCGTCAACCACAAAAATATAATTGGCTTACTAAACGTGTTCACACCACAG
AAGACCCTTGAAGAGTTCCAGGATGTTTATCTGGTGATGGAGCTGATGGATGCAAAC
CTGTGTCAGGTGATTCAGATGGAGCTGGACCACGAGAGGCTGTCCTACCTGCTCTAT
CAGATGCTCTGCGGCATTAAACACCTGCACGCTGCTGGCATCATACACAGGGACCTG
AAACCCAGTAATATAGTAGTGAAATCGGACTGCACGCTGAAGATCCTGGATTTCGGT
CTGGCCAGAACGGCTGCAACCGGCCTCCTCATGACTCCTTATGTAGTGACACGCTAT
TATCGGGCCCCAGAGGTCATCCTGGGCATGGGTTATCAAGCTAACGTTGATGTCTGG
TCTATTGGCTGCATCATGGCTGAAATGGTCAGAGGTAGTGTGTTGTTTCCTGGCACA
GACCATATTGACCAGTGGAATAAAGTGATCGAGCAGCTCGGCACGCCGTCACAGGAG
TTCATGATGAAGCTGAATCAGTCTGTGAGGACGTATGTGGAGAACCGGCCTCGGTAT
GCGGGATACAGCTTTGAGAAGCTCTTCCCAGACGTGCTCTTCCCCGCAGACTCGGAC
CACAACAAACTCAAGGCGAGTCAGGCACGAGACTTGTTATCCAAAATGCTGGTAATA
GATGCGTCCAAGCGGATCTCTGTAGACGAGGCGCTTCAGCACCCCTACATCAACGTT
TGGTACGACCCGTCAGAAGTGGAGGCGCCACCACCAGCGATCACGGATAAACAGCTC
GATGAGAGAGAACACTCAGTGGAAGAGTGGAAAGAGCTCATATATAAGGAAGTGCTG
GAATGGGAGGAGCGAACAAAAAATGGAGTGATCAGAGGACAGCCGGCCTCGCTAGCA
CAGGTGCAGCAGTGA

> jnk1b_ex8_sh Protein
MNRNKREKEYYSIDVGDSTFTVLKRYQNLRPIGSGAQGIVCSAYDHVLDRNVAIKKL
SRPFQNQTHAKRAYRELVLMKCVNHKNIIGLLNVFTPQKTLEEFQDVYLVMELMDAN
LCQVIQMELDHERLSYLLYQMLCGIKHLHAAGIIHRDLKPSNIVVKSDCTLKILDFG
LARTAATGLLMTPYVVTRYYRAPEVILGMGYQANVDVWSIGCIMAEMVRGSVLFPGT
DHIDQWNKVIEQLGTPSQEFMMKLNQSVRTYVENRPRYAGYSFEKLFPDVLFPADSD
HNKLKASQARDLLSKMLVIDASKRISVDEALQHPYINVWYDPSEVEAPPPAITDKQL
DEREHSVEEWKELIYKEVLEWEERTKNGVIRGQPASLAQVQQ*
```

>jnk1b_ex8_Lg
ATGAACAGGAATAAGCGCGAGAAAGAATATTACAGCATAGATGTAGGAGATTCGACG
TTCACCGTTTTGAAGCGCTATCAGAATTTAAGACCAATCGGGTCCGGAGCACAAGGC
ATCGTCTGCTCAGCGTATGACCACGTCCTCGATCGAAATGTGGCGATTAAGAAACTC
AGCCGACCCTTTCAAAACCAAACTCATGCCAAACGGGCCTACAGAGAACTGGTCCTG
ATGAAATGCGTCAACCACAAAAATATAATTGGCTTACTAAACGTGTTCACACCACAG
AAGACCCTTGAAGAGTTCCAGGATGTTTATCTGGTGATGGAGCTGATGGATGCAAAC
CTGTGTCAGGTGATTCAGATGGAGCTGGACCACGAGAGGCTGTCCTACCTGCTCTAT
CAGATGCTCTGCGGCATTAAACACCTGCACGCTGCTGGCATCATACACAGGGACCTG
AAACCCAGTAATATAGTAGTGAAATCGGACTGCACGCTGAAGATCCTGGATTTCGGT
CTGGCCAGAACGGCTGCAACCGGCCTCCTCATGACTCCTTATGTAGTGACACGCTAT
TATCGGGCCCCAGAGGTCATCCTGGGCATGGGTTATCAAGCTAACGTTGATGTCTGG
TCTATTGGCTGCATCATGGCTGAAATGGTCAGAGGTAGTGTGTTGTTTCCTGGCACA
GACCATATTGACCAGTGGAATAAAGTGATCGAGCAGCTCGGCACGCCGTCACAGGAG
TTCATGATGAAGCTGAATCAGTCTGTGAGGACGTATGTGGAGAACCGGCCTCGGTAT
GCGGGATACAGCTTTGAGAAGCTCTTCCCAGACGTGCTCTTCCCCGCAGACTCGGAC
CACAACAAACTCAAGGCGAGTCAGGCACGAGACTTGTTATCCAAAATGCTGGTAATA
GATGCGTCCAAGCGGATCTCTGTAGACGAGGCGCTTCAGCACCCCTACATCAACGTT
TGGTACGACCCGTCAGAAGTGGAGGCGCCACCACCAGCGATCACGGATAAACAGCTC
GATGAGAGAGAACACTCAGTGGAAGAGTGGAAAGAGCTCATATATAAGGAAGTGCTG
GAATGGGAGGAGCGAACAAAAAATGGAGTGATCAGAGGACAGCCGGCCTCGCTAGGT
GCAGCAGTGAGCAGTGACTCCCATGAGCCCTCGACGTCGTCCTCCTCCATAAACGAT
GTGTCGTCCATGTCCACCGAGGTCACGCTGACCTCAGACACCGACAGCAGTCAGGAG
ACGTCCAACGGAGCGCTGCACTGCTGCAGATGA

> jnk1b_ex8_Lg Protein
MNRNKREKEYYSIDVGDSTFTVLKRYQNLRPIGSGAQGIVCSAYDHVLDRNVAIKKL
SRPFQNQTHAKRAYRELVLMKCVNHKNIIGLLNVFTPQKTLEEFQDVYLVMELMDAN
LCQVIQMELDHERLSYLLYQMLCGIKHLHAAGIIHRDLKPSNIVVKSDCTLKILDFG
LARTAATGLLMTPYVVTRYYRAPEVILGMGYQANVDVWSIGCIMAEMVRGSVLFPGT
DHIDQWNKVIEQLGTPSQEFMMKLNQSVRTYVENRPRYAGYSFEKLFPDVLFPADSD
HNKLKASQARDLLSKMLVIDASKRISVDEALQHPYINVWYDPSEVEAPPPAITDKQL
DEREHSVEEWKELIYKEVLEWEERTKNGVIRGQPASLGAAVSSDSHEPSTSSSSIND
VSSMSTEVTLTSDTDSSQETSNGALHCCR*
